# Genotype inference from aggregated chromatin accessibility data reveals genetic regulatory mechanisms

**DOI:** 10.1101/2024.09.04.610850

**Authors:** Brandon M. Wenz, Yuan He, Nae-Chyun Chen, Joseph K. Pickrell, Jeremiah H. Li, Max F. Dudek, Taibo Li, Rebecca Keener, Benjamin F. Voight, Christopher D. Brown, Alexis Battle

**Author notes:** These authors contributed jointly to the work (Brandon M. Wenz, Yuan He). Correspondence to: Alexis Battle, PhD.

## Abstract

**Background:** Understanding the genetic causes for variability in chromatin accessibility can shed light on the molecular mechanisms through which genetic variants may affect complex traits. Thousands of ATAC-seq samples have been collected that hold information about chromatin accessibility across diverse cell types and contexts, but most of these are not paired with genetic information and come from diverse distinct projects and laboratories.

**Results:** We report here joint genotyping, chromatin accessibility peak calling, and discovery of quantitative trait loci which influence chromatin accessibility (caQTLs), demonstrating the capability of performing caQTL analysis on a large scale in a diverse sample set without pre-existing genotype information. Using 10,293 profiling samples representing 1,454 unique donor individuals across 653 studies from public databases, we catalog 23,381 caQTLs in total. After joint discovery analysis, we cluster samples based on accessible chromatin profiles to identify context-specific caQTLs. We find that caQTLs are strongly enriched for annotations of gene regulatory elements across diverse cell types and tissues and are often strongly linked with genetic variation associated with changes in expression (eQTLs), indicating that caQTLs can mediate genetic effects on gene expression. We demonstrate sharing of causal variants for chromatin accessibility and diverse complex human traits, enabling a more complete picture of the genetic mechanisms underlying complex human phenotypes.

**Conclusions:** Our work provides a proof of principle for caQTL calling from previously ungenotyped samples, and represents one of the largest, most diverse caQTL resources currently available, informing mechanisms of genetic regulation of gene expression and contribution to disease.

## Introduction

Genome wide association studies (GWAS) have identified thousands of loci and common human genetic variants that are associated with a wide range of complex human traits, diseases, and risk factors[1]. GWAS variants are often found in noncoding regions, where they are likely to be involved in gene regulation[2,3]. However, a full picture of the causal regulatory elements that underlie these associations remains incomplete for most loci[4]. Characterizing the genetic effects of variants on gene expression as revealed by expression quantitative trait locus (eQTL) mapping has provided insights into the molecular basis of phenotypes[3,5–7].

Although some eQTL variants directly affect open-reading frames, the vast majority are in non-coding regions, as has been described for GWAS variants. Connecting causal variants to the regulatory elements and the genes of action that they perturb remains a central goal of the post-GWAS era.

Accessibility of chromatin regions to transcriptional machinery is a key factor in gene regulation[8,9], and genetic variants can affect complex traits through changes in gene expression levels that are mediated by chromatin accessibility[10,11]. Improved understanding of the mechanisms involved in chromatin accessibility, revealed by genetic variants that modulate chromatin accessibility (i.e., caQTLs), has the potential to illuminate the molecular mechanisms and genetic regulatory architecture of complex traits. caQTLs have been measured in a variety of tissue and cell types, at both bulk[12–16] and single-cell resolutions[17]. caQTLs have been used in a variety of studies to characterize gene expression regulation[18], and to propose mechanisms for risk loci identified through GWAS[19]. caQTLs may co-occur with eQTLs together, thus describing a more complete picture of the genetic mechanism underlying GWAS-associated signals. However, relevant caQTLs may be discovered even in the absence of any established eQTL, as eQTL studies may not include the relevant cell type or environmental context to reveal the change to gene expression. Analysis of the contribution of caQTLs to complex human traits can help us better understand the molecular impact of these variants and the mechanism(s) driving GWAS signals. To date, caQTL studies have mostly been performed in analyses restricted to single tissue/cell types, a majority of which have assayed a limited number of samples.

The Assay for Transposase-Accessible Chromatin using sequencing (ATAC-seq) technology has been widely used to capture chromatin accessibility in various cell types and experimental conditions[20–22]. There is a rapidly accumulating trove of ATAC-seq data generated from various experiments, labs, and conditions. This wealth of information has the potential to boost power for caQTL analysis. Unfortunately, many of these samples do not have matched genotype information, a necessary component for QTL analyses. ATAC-seq reads, however, naturally carry the sequence information at nucleotide resolution, providing the possibility of inferring sample genotypes from these data directly.

Here, we have selected and evaluated pipelines to uniformly process ATAC-seq samples, including peak calling and genetic variant calling directly from ATAC-seq reads. We called genotypes using a pipeline incorporating Gencove’s low-pass sequencing methods applied to ATAC-seq reads in accessible chromatin, which utilizes imputation to infer genotype for variants that are located outside of regions covered by observed reads in accessible regions[23,24]. We benchmarked this pipeline, using gold standard genotype information available for a subset of samples, and compared it with other methods. Because large-scale public data often contains multiple samples from the same donor or even the same cell line, we also developed a method to automatically infer donor assignment based on genotype from the called variants. Peak calling from thousands of diverse samples presents challenges of identifying true, distinct regions of chromatin accessibility rather than low-signal false positives, or large regions merged from what should be distinct peaks[25,26]. Based on comparisons across various peak-calling approaches, we finalized a pipeline based on an Genrich, an ATAC-seq specific method[27] for collectively calling peaks across large, diverse data sets and quantifying accessibility in each peak.

Using our ATAC-seq derived genotypes and accessibility estimates across peaks and samples, we then called caQTLs from this collection of publicly available ATAC-seq data. We identified thousands of caQTLs that share a causal signal with GWAS signals, many of which are not explained by known eQTLs. Additionally, we identified many GWAS signals that appear to share a causal signal with both eQTLs and caQTLs, enabling a more comprehensive analysis predicting target gene, gene regulatory element and even potential transcription factors that are driving GWAS signals for a variety of complex human traits. Furthermore, to capture context-specific caQTLs, we inferred clusters of samples with similar accessibility profiles, mostly reflecting cell or tissue type, and identified cluster-specific caQTLs. With the captured global and cluster-specific caQTLs, we investigated potential mechanisms involving transcription factors and their role in target gene regulation.

## Results

### Accurate genotyping and imputation based on ATAC-seq reads from public repositories

We established a workflow to collect a diverse set of publicly available ATAC-seq datasets and ascertain donor genotype from ATAC-seq reads, with the overall objective of mapping genetic variants that are associated with differences in chromatin accessibility for diverse tissues and contexts on a large scale (Figure 1A). We collected 10,293 human samples from 653 projects from the Gene Expression Omnibus (GEO) data repository, where most projects were comprised of 10 or fewer samples (Figure 1B, Supplementary Table 1). The aggregated data includes samples from a wide variety of tissues or cell types (Figure 1C), labeled based on a manual curation of project abstracts, sample labels, and project methods, with the most common cell/tissue types including T cells and brain. Additionally, based on our metadata review, both cancer and normal primary tissue are well represented, along with cell lines and experimentally differentiated cell types (Figure 1D). The diversity of samples highlights the value of a workflow that can aggregate data and genotype samples from ATAC-seq reads, providing an overall large sample size, but also tissue-specific sample sizes larger than any existing genotyped chromatin accessibility study for several individual tissues including lung, breast, heart, and pancreas[12,28–30].

**Figure 1.**
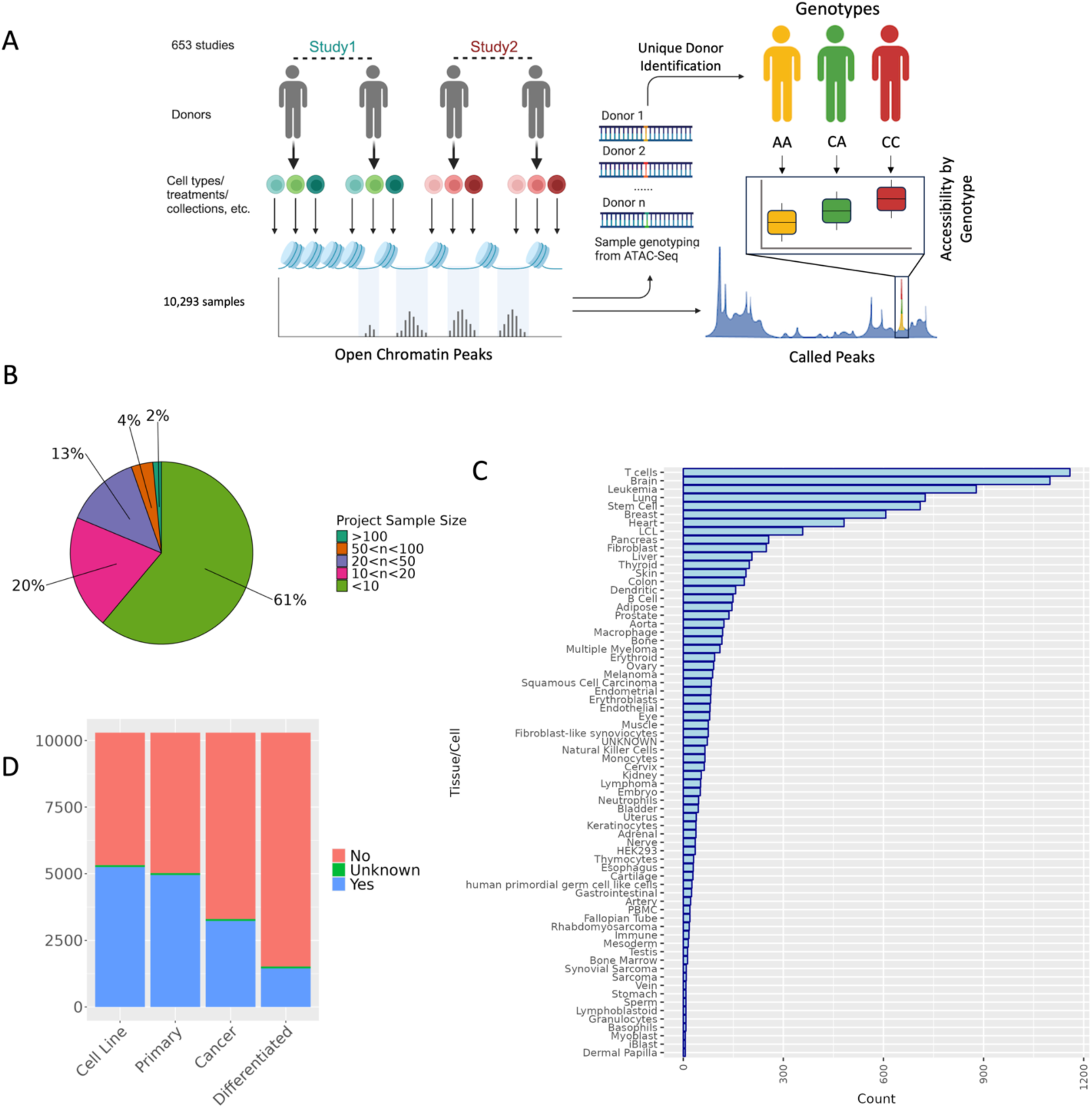
Study overview and characteristics of specimens utilized in this study. **(A)** Overview of study design to jointly call genotype and caQTLs across studies. Human ATAC-seq datasets were obtained from GEO. After variant-calling (Methods), we identified the unique donors in the dataset (Methods) for use in caQTL mapping. **(B)** The distribution of the number of samples collected across all n=653 studies. **(C)** Frequency of the Cell/Tissue types present in samples collected across studies based on manual metadata curation **(D)** Frequencies of cancer, non-cancer, primary tissues, and cell-line samples included in our study based on our metadata review. For each category, samples were assigned a “Yes” if they belonged to that category (e.g. cell line samples for ‘Cell Line’ category), a “No” if they did not belong (e.g. primary tissue samples for ‘Cell Line’ category), or an “Unknown” if it was not clear from the metadata.

QTL mapping requires paired genotype and molecular phenotype information for each sample. In standard QTL studies, genotyping arrays or whole genome sequencing (WGS) are used to ascertain sample genotype information[31]. Unfortunately, for most of the ATAC-seq data in public repositories that has already been collected, genotype data is not readily available. However, ATAC-Seq directly captures genomic DNA fragments from accessible chromatin regions; thus, we surmised that it might instead be possible extract genotype information for these samples directly from the ATAC-seq reads. To obtain genotyping from ATAC-sequencing and evaluate the performance of variant calling using ATAC-seq reads, we applied two approaches: a pipeline incorporating genotyping from Gencove, which optimizes genotyping and imputation for low-pass sequencing data[23,24,32,33], and a standard GATK variant calling pipeline[32,33](Methods). To benchmark the performance of our workflow, we used a published dataset of 71 HapMap lymphoblastoid cell lines (LCL) samples with paired ATAC-seq and WGS data [34]. We observed that, compared to the standard GATK variant calling pipeline, the Gencove pipeline with imputation greatly increased the number of variants called and resulted in a median correlation of over 0.88 between true and called donor genotype (Figure 2A). To quantify the effects of read coverage on the performance of variant calling, we randomly subselected ATAC-seq reads at varying total read counts for use with the Gencove pipeline. We observed a marginal increase in accuracy with deeper coverage, however, variant-calling accuracy remained high at effective coverage as low as 0.04 (Figure 2B). In our full dataset, the distribution of effective coverage in the full sample set was within the range previously tested with the gold standard HapMap LCL samples, verifying the accuracy of genotype calling in this larger data set. These analyses demonstrate the capabilities of accurate inference of genome-wide genotypes directly from ATAC-seq data.

**Figure 2.**
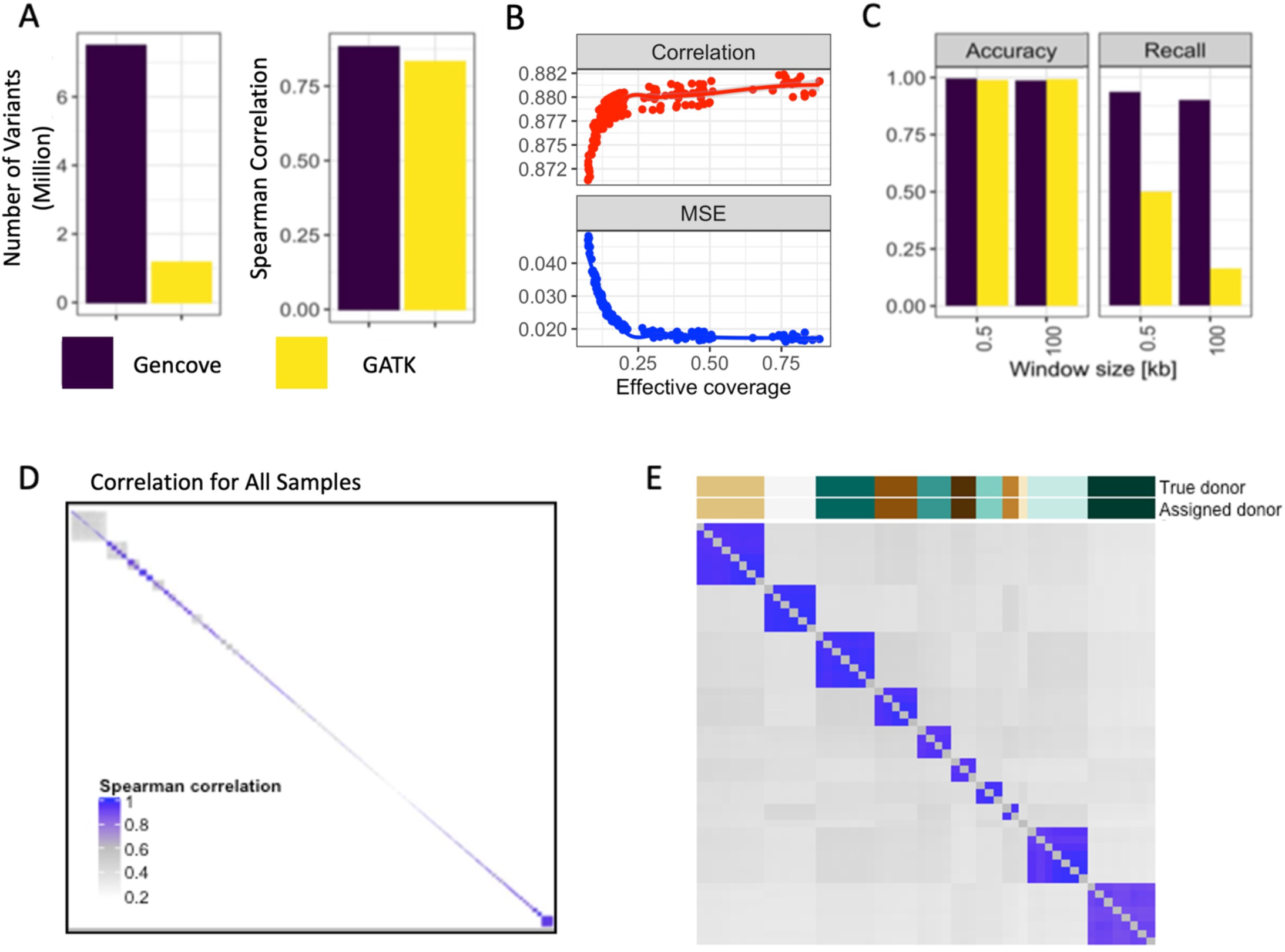
High quality genotyping with unique donor information is inferable directly from reads obtained by ATAC-Seq. **(A)** Variants called for the HapMap samples using two pipelines - Gencove, and GATK HaplotypeCaller. **(B)** Accuracy of variant genotype called by Gencove pipeline using a random subset of sample reads. Spearman correlation and mean squared error (MSE) are computed between the called genotype and genotype from WGS. **(C)** caQTLs called using ATAC-seq derived genotypes across the HapMap samples. **(D)** Spearman correlation of called genotypes between all samples. **(E)** Spearman correlation of called genotypes between samples in study PRJNA388006. On the top the “True donor” indicates the donor assignment obtained from metadata information for this study, and “Assigned donor” indicates the donor assignment derived from called genotypes **(Methods)**.

As a proof of concept, we next performed caQTL mapping using genotypes called from ATAC-seq reads, comparing the results to the caQTLs identified using the full set of gold standard genotypes in these 71 HapMap LCL samples. We observed that caQTL calling using ATAC-seq reads and the Gencove pipeline performed better than the GATK pipeline with 99% accuracy and over 90% recall compared to caQTL calling using WGS data. The increased recall is due to the Gencove pipeline’s imputation step and sacrifices very little in accuracy (Figure 2C). The performance of the Gencove pipeline had substantially greater benefit when testing variants in larger caQTL mapping window sizes where recall remained above 90% for the Gencove pipeline but dropped to 16% for the GATK pipeline at 100 kb (Figure 2C). Overall, we conclude that genotype calling from ATAC-seq reads leads to highly accurate caQTL calling with relatively high recall with a low rate of false positives. Given the diverse samples collected and varying study designs, an individual donor will likely have multiple ATAC-seq samples represented. As such, we next developed a pipeline to infer unique donors based on the correlation between inferred sample genotypes across different samples and projects (Figure 2D-E, Methods). Applying this pipeline to all samples, we identified 1,454 unique donors across our entire dataset (Supplementary Table 2). The majority of donors (∼82%) are found within a single project only. As expected, the occurrence of multiple samples per donor was especially common amongst cell lines, which is reflected in the reduced proportion of cell line samples in the final unique donor sample set (Supplementary Figure 1).

### Peak calling across all samples identifies a plethora of open chromatin regions with regulatory potential

The next step in our pipeline was to identify open chromatin regions. We called chromatin accessibility peaks based on evidence across all samples using Genrich, a peak caller optimized for ATAC-seq reads[27]. Genrich assigns p-values to genomic positions within each sample, then combines p-values across samples using Fisher’s method to call peaks. We compared this Genrich pipeline to strategies which called peaks in individual samples followed by merging. The Genrich strategy alone produced peaks that are likely derived from nucleosome-free and mono-nucleosome fragments, as seen by enrichment around 100 bp and 200 bp in the observed peak length distribution (Figure 3A).

**Figure 3.**
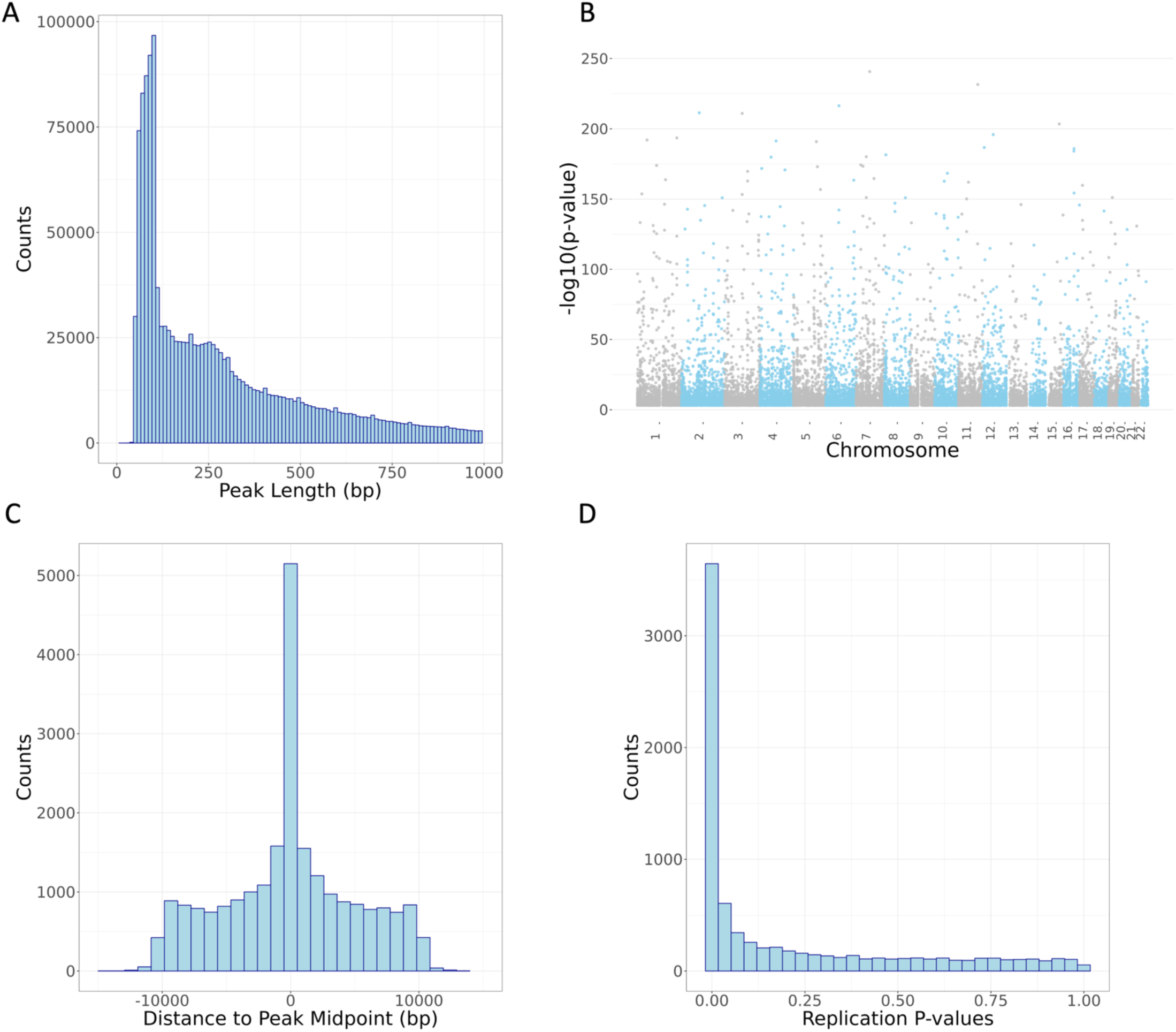
Characteristics of chromatin accessibility peaks and caQTL variants identified in this study. **(A)** Distribution of peak length across 1,659,379 called peaks (peaks under 1000 bp shown). **(B)** Manhattan plot of lead variant for 23,381 caQTL peaks. **(C)** Distance from lead caQTL variant to midpoint of caQTL peak showing elevation of caQTL variant within the identified chromatin accessibility peak. **(D)** Lead variants for 23,381 caQTL peaks were matched in external caQTL mapping dataset of African LCLs[49]; p-values from the replication study are plotted here.

Across 10,293 samples, we identified 1,659,379 autosomal peaks with a median peak length of 250 base pairs, covering approximately 27% of the genome (Figure 3A). Chromatin accessibility is influenced by a variety of regulatory processes[35–37], and we would expect to see chromatin accessibility peaks in regions associated with gene regulation. To verify the quality of our ATAC-seq peaks, we annotated our peaks, along with length-matched, randomly selected controls, with various genomic features that included transcript annotations and enhancer annotations as defined by the FANTOM5 enhancer atlas[38,39] (Methods). We found that relative to controls, our ATAC-seq peaks were enriched for genomic regions annotated as enhancers and all transcript annotations but depleted for gene intergenic regions (Supplementary Figure 2, Supplementary Table 3). Similarly, we would expect our ATAC-seq peaks to be enriched for histone modifications associated with gene regulatory regions[40–42]. The ENCODE Roadmap Epigenomics Mapping Consortium[43] provides chromatin immunoprecipitation with sequencing (ChIP-seq) data representing eight different histone marks from 556 cell line, tissue, and primary cell samples derived from a variety of biological origins. Using these data, the highest enrichment of our ATAC-seq peaks and chromatin histone marks was for H3K4me1, a histone mark that has been linked to enhancers (Supplementary Table 4)[40]. In contrast, our ATAC-seq peaks were depleted for overlap with the histone mark H3K9me3, which is associated with gene repression and heterochromatin[44]. Together, these data suggest that our ATAC-seq peaks are enriched for cis-regulatory regions, as expected for genomic sequences implicated in regulatory activity and indicating high quality peak calls.

### Inferred genotypes support high-powered caQTL mapping across samples

Next, we sought to identify genetic variants that are associated with differences in measured chromatin accessibility in ATAC-seq peaks, i.e., caQTLs. We tested a 10 kilobase (kb) window in *cis* flanking each chromatin accessibility peak, as we anticipate that genetically altered active transcription factor binding sites are likely to be found within or very nearby regions of chromatin accessibility[45,46]. Utilizing our peak calling and genotyping pipelines, we identified 23,381 chromatin accessibility peaks with a significant caQTL at FDR 5% across 1,454 unique donor samples (Figure 3B, Methods, Supplementary Tables 5-6). To mitigate potential confounding from population stratification, we estimated variation in similarity across donors generated by our genotyping via principal components analysis (PCA), including 3 PCs as covariates in discovery analysis. In addition, we also included 200 PCs generated from the donor chromatin accessibility peak read count matrix to mitigate potential latent confounders in QTL mapping [47] (Methods).

We examined the quality of our caQTL variants by determining whether they were enriched for expected functional characteristics. First, we confirmed that the distribution of positions for lead caQTL variants was centered within the open chromatin peak tested, as expected (Figure 3C). In addition, we observed that peaks with a mapped caQTL were the most strongly enriched for gene 5’ UTRs and enhancer regions while depleted in gene intergenic regions (Supplementary Figure 3, Supplementary Table 7). Interestingly, caQTL peaks were further enriched in enhancer regions compared to all chromatin accessibility peaks, suggesting that caQTLs we mapped may be found at genomic elements involved in distal gene regulation. This could potentially arise due to selective pressure reducing functional variation in promoters and other proximal elements.

Additionally, we examined whether our caQTL peaks were enriched for transcription factor binding sites in the ENCODE transcription factor ChIP-seq data from 129 cell types and 340 transcription factors[48]. As expected, caQTL peaks, compared to length-matched random controls, were enriched for binding sites for all transcription factors except for SRSF9, which is depleted in caQTL peaks (Supplementary Table 8). Enrichment of these functional characteristics support the conclusion that our caQTLs are high quality, reflect enrichment in expected regulatory elements, and can help identify genetic mechanisms relevant to regulation of gene expression. We sought further evidence that caQTL variants were enriched for functional roles in gene expression regulation by intersecting them with eQTLs. Across all 49 Genotype-Tissue Expression (GTEx) v8 tissues, we observed caQTL/eQTL enrichments ranging from 2.1 to 4.8-fold per tissue and a total of 2,859 (∼13% of unique caQTLs) unique overlapping lead caQTL/lead eQTL variants found across all tissues, for an enrichment of approximately 1.8-fold (Supplementary Table 9).

Finally, to further demonstrate that our catalog represents reproducible peaks and caQTLs, we compared our findings here to a recent caQTL study that identified variants associated with chromatin accessibility in African LCL samples[49] not included in our discovery effort. Lead caQTLs and peaks identified in our study resulted in a replication rate (π_1_ value[50,51]) of 0.62 with this orthogonal study (Figure 3D). Together, these analyses further demonstrate that on average, our catalog of caQTLs are high quality and provide insight into how genetic variation may affect gene regulation and complex traits.

### Colocalization suggests shared causality between chromatin accessibility, complex traits, and expression QTLs

To gain further insight into the molecular mechanisms underlying GWAS signals, we sought to link GWAS association signals, expression QTLs (eQTLs), and caQTLs together via statistical colocalization (**Methods**). Colocalization analysis discerns if an association signal is likely shared between two traits, suggestive of a common underlying genetic mechanism. First, we examined which caQTL signals are shared with GWAS signals across a variety of complex human traits. We obtained GWAS summary statistics from a subset of the UK Biobank (UKBB) study, selecting 78 traits of interest with high confidence of significant heritability (**Methods**)[52]. We then performed colocalization analysis (**Methods**) for any caQTL peak that was located within 1 Mb of a genome-wide significant lead GWAS signal (**Methods**). We observed that 67 traits had a caQTL/GWAS colocalization event (PP3+PP4 > 0.8 and PP4/(PP3+PP4) > 0.9.) for a total of 12,882 colocalization events across all traits, involving 4,351 (∼19%) unique caQTL peaks and 4,706 (∼34%) unique tested GWAS signals (Supplementary Table 10).

Regulatory variants do not always affect the nearest gene and assigning a GWAS signal to a causal gene is not a trivial procedure[53,54]. Furthermore, comparison of the overlap between lead variants of GWAS signals and the lead variant of eQTLs can suggest the incorrect causal gene[55]. Given the prominence of long-range gene expression regulation, colocalization of cis regulatory elements with eGenes can suggest a shared causal variant[56,57]. We performed colocalization analyses between caQTLs and 49 GTEx v8 eQTL tissues. Across all tissues, between 358 (Kidney) and 5,427 (Thyroid) eGenes colocalized with our caQTLs.

Colocalized caQTLs/eQTLs were shared across a median of three tissues and 17,471 unique eGenes colocalized with caQTLs in any GTEx tissue (Supplementary Figure 4, Supplementary Table 11). We found that only 13% of eQTL/caQTL colocalizations involve the gene nearest to the lead caQTL and that there was a median of 6 genes closer to the lead caQTL than the colocalizing gene (Supplementary Figure 4). Additionally, the putative regulated gene transcription start site (TSS) was a median of 80,798 base pairs away from the colocalizing caQTL (Supplementary Figure 4). These results suggest that caQTLs may often be found tagging and potentially modifying the behavior of distal gene regulatory elements.

### Multiple molecular QTL datasets provide insight into regulatory mechanisms underlying GWAS associations

eQTLs have been shown to provide a regulatory mechanistic hypothesis for GWAS associated signals, yet only an estimated ∼25-43% of GWAS signals colocalize with known eQTLs[6,58], implying that more than half of GWAS loci may lack an obvious functional, mechanistic hypothesis[6,59–61]. caQTL mapping could help close that gap if, for example, the effects of the eQTL are only apparent in certain cellular contexts, during specific developmental stages, or in the presence of external stimuli[62–64], whereas chromatin accessibility may be primed and reveal effects in a wider range of context. Across all traits and GTEx tissues, we find that lead GWAS signals colocalize with a median of 5 eQTLs and 2 caQTLs (Supplementary Figure 5). For each GWAS trait, we then considered whether independent GWAS lead signals colocalize only with eQTLs, colocalize with both caQTLs and eQTLs, or colocalize only with caQTLs. Across all GWAS, a median of 35 unique signals colocalized with a caQTL only, a median of 66 unique signals colocalized with an eQTL only, and a median of 53 unique signals colocalized with both a caQTL and an eQTL (Figure 4, Supplementary Table 13). These differences may reflect context-specific behavior of gene regulation that is not well captured by steady-state, adult gene expression data, but may still be reflected in chromatin accessibility. These results demonstrate that incorporating both caQTLs and eQTLs nominates putative causal mechanisms for approximately 29% more GWAS signals than using eQTLs alone.

**Figure 4.**
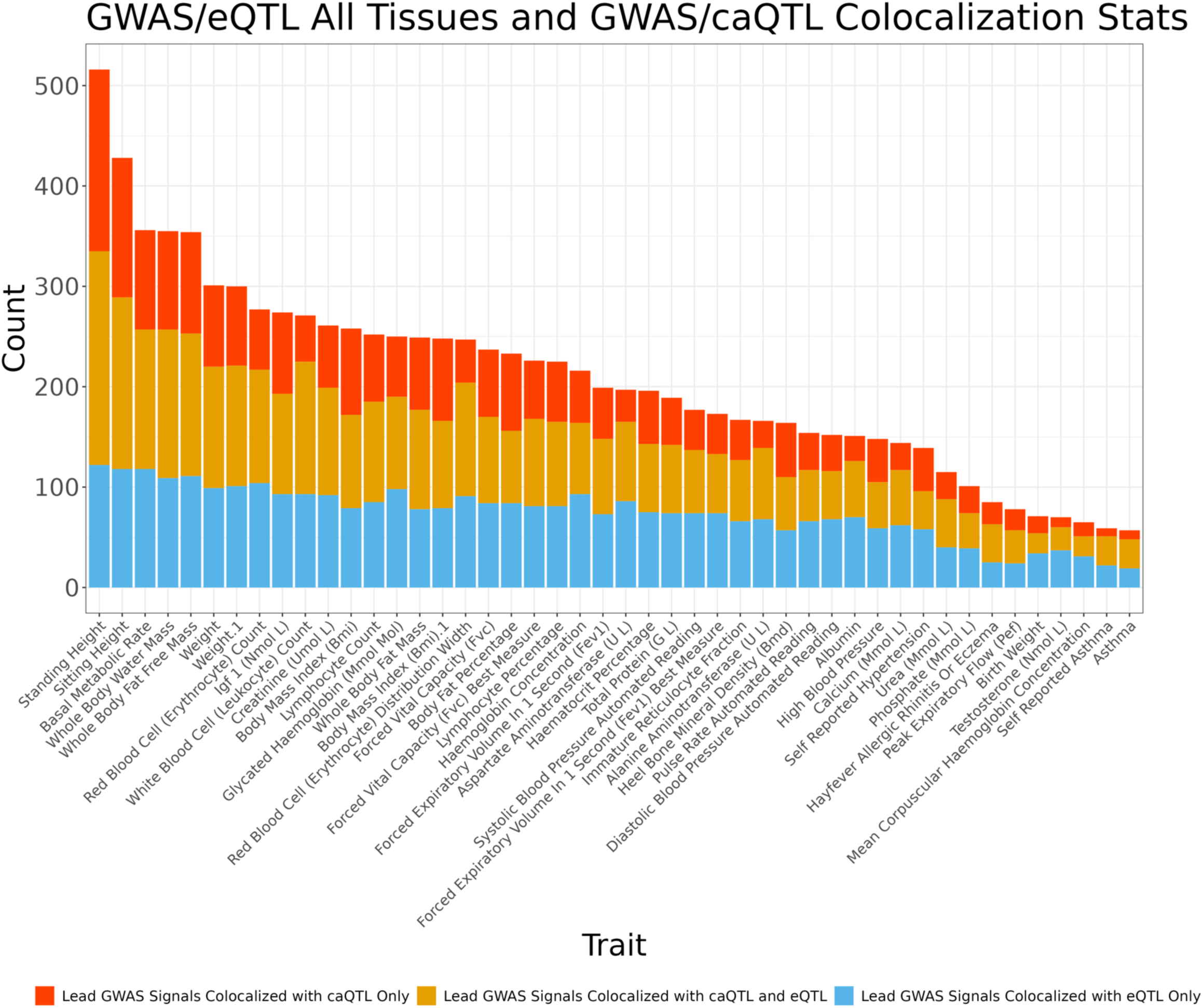
caQTLs map to regions tagged by GWAS and eQTL variation. For each GWAS trait, independent lead GWAS variant signals were checked for colocalization with caQTL and eQTL signals across all GTEx tissues. Plotted is the number of unique lead GWAS signals per colocalization group, as multiple caQTL peaks, eGenes, etc. can colocalize with the same lead GWAS signal. Traits with greater than 50 colocalizing lead variants shown.

Furthermore, 57% of GWAS signals we tested were linked with either a caQTL, eQTL, or both (Supplementary Figure 6). Instances where GWAS signals colocalized with both caQTLs and eQTLs may also allow for a better delineation of the mechanism at these loci by nominating a candidate caQTL-associated gene regulatory element to a target eGene[65].

To gain insight into molecular mechanisms that may be unique to caQTLs as compared to eQTLs, we calculated the enrichment of colocalizing caQTLs and lead eQTLs for diverse genomic annotations. caQTLs and eQTLs involved in colocalizations with GWAS signals were both significantly enriched for all tested genomic annotation categories except for intergenic regions, where they were significantly depleted, compared to matched random controls (Supplementary Figure 7, **Methods**). However, caQTLs from GWAS/caQTL and caQTL/GWAS/eQTL colocalization events were further enriched for enhancer regions and less depleted in intergenic regions than eQTLs from GWAS/eQTL colocalizations alone (Supplementary Figures 8-9). In contrast, lead variants of eQTLs that colocalized with a GWAS signal only were further enriched for gene promoters and other gene proximal categories, less enriched in enhancer regions, and showed greater depletion for intergenic regions, consistent with previous reports (Supplementary Figure 10)[6,66]. These differences in enrichment may be due to systematic differences in GWAS signals that are explained by eQTLs compared to those explained by potentially distal regulatory mechanisms captured by caQTLs[67].

While our caQTLs were called from heterogeneous cell/tissue samples, they are predominantly from brain and whole blood (Figure 1). To reflect this, we also performed an analysis of caQTL/GWAS colocalizations compared to eQTL/GWAS colocalizations from brain cortex and whole blood only. Across 69 GWAS, each trait has at least 1 GWAS signal that colocalizes only with a caQTL, and one trait, standing height, had 360 lead GWAS variants that colocalize exclusively with caQTLs compared to brain eQTLs. In contrast, we identify a maximum of 66 lead GWAS variants that colocalize only with eQTLs for a given trait. Across all GWAS, a median of 76 unique signals colocalized with a caQTL only, a median of 15 unique signals colocalized with an eQTL only in Whole Blood, and a median of 11 unique signals colocalized with both a caQTL and a Whole Blood eQTL (Supplementary Figure 11, Supplementary Table 14). Furthermore, across all GWAS, a median of 83 unique signals colocalized with a caQTL only, a median of 10 unique signals colocalized with an eQTL only in Brain Cortex, and a median of 7 unique signals colocalized with both a caQTL and a Brain Cortex eQTL (Supplementary Figure 12, Supplementary Table 15). Compared to the analysis considering eQTLs across all tissues, we find that caQTL/GWAS only colocalizations occur with a larger proportion of GWAS signals in single tissue eQTL analysis colocalizations. This discrepancy provides further evidence that using caQTLs can provide molecular insight into GWAS association signals beyond eQTLs when restricting to a single eQTL tissue.

### Integration of caQTLs informs mechanistic interpretation at many GWAS loci

Colocalization analysis with QTL datasets across multiple modalities, such as expression and chromatin accessibility, has previously been shown to nominate putative target genes underlying more GWAS signals than a single modality alone[65,68]signals that colocalized separately with both caQTLs and eQTLs and quantified how many of the GWAS-colocalizing caQTLs and eQTLs also colocalized with each other. We identified 43,005 unique colocalization events involving a GWAS trait, caQTL peak, eGene, and eGene tissue (Supplementary Table 16). These were comprised of 2,177 unique eGenes and 1,695 unique caQTL peaks.

In cases where caQTLs colocalize with both GWAS signals and eQTLs, they provide a more complete picture of the mechanisms likely driving the association signal. First, we provide an instructive example of a well-characterized GWAS locus strongly associated with plasma low-density lipoprotein cholesterol (LDL-C) at the 1p13 locus. eQTL colocalization analyses at this locus, followed by functional characterization in vitro and in vivo, suggest that the causal gene at this locus is *SORT1*, with expression differences observed in the liver [46]. We find a caQTL at this locus that colocalizes with both the *SORT1* eQTL in liver, and the GWAS trait self-reported high cholesterol (Supplementary Figure 13). This caQTL peak contains a well-studied noncoding variant that creates a C/EBP (CCAAT/enhancer binding protein) TF binding site, altering hepatic expression of *SORT1* and plasma LDL-C levels[46]. This highlights the ability of our analyses to identify verified mechanisms underlying GWAS signals.

In a second example, we identified a compelling locus where a caQTL peak, a whole blood eQTL for *PAX8*, and a GWAS signal for blood urea levels colocalized (Figure 5). The shared lead caQTL and eQTL variant, rs7589901, is an intronic variant within the *PAX8* gene. The reference allele of rs7589901-A is associated with increased chromatin accessibility in the associated peak (Supplementary Figure 14). Based on motif analysis, ZNF135 is predicted to bind to a motif overlapping rs7589901, with the alternate C allele strongly favored for binding (PWM value=0.8, Supplementary Figure 15). In GTEx, the rs7589901 eQTL direction of effect is concordant with the caQTL direction of effect, suggesting that increased accessibility at this locus is associated with increased *PAX8* gene expression in whole blood. The lead GWAS variant at this locus, rs7421852, is associated with increased blood urea levels, is ∼3,000 bp from rs7589901, and is in strong LD (r^2^=0.85) with rs7589901 in our caQTL sample genotypes. These results suggest a potential mechanism where ZNF135 is acting as a transcriptional repressor at this locus, a functional role that has been implicated in a different context[69]. The culmination of evidence suggests a mechanism where decreased ZNF135 binding leads to increased chromatin accessibility, increased expression of the *PAX8* gene, and lower blood urea levels. Such examples demonstrate the power of integrating multiple molecular QTL datasets to nominate mechanistic hypotheses that may be further validated experimentally.

**Figure 5.**
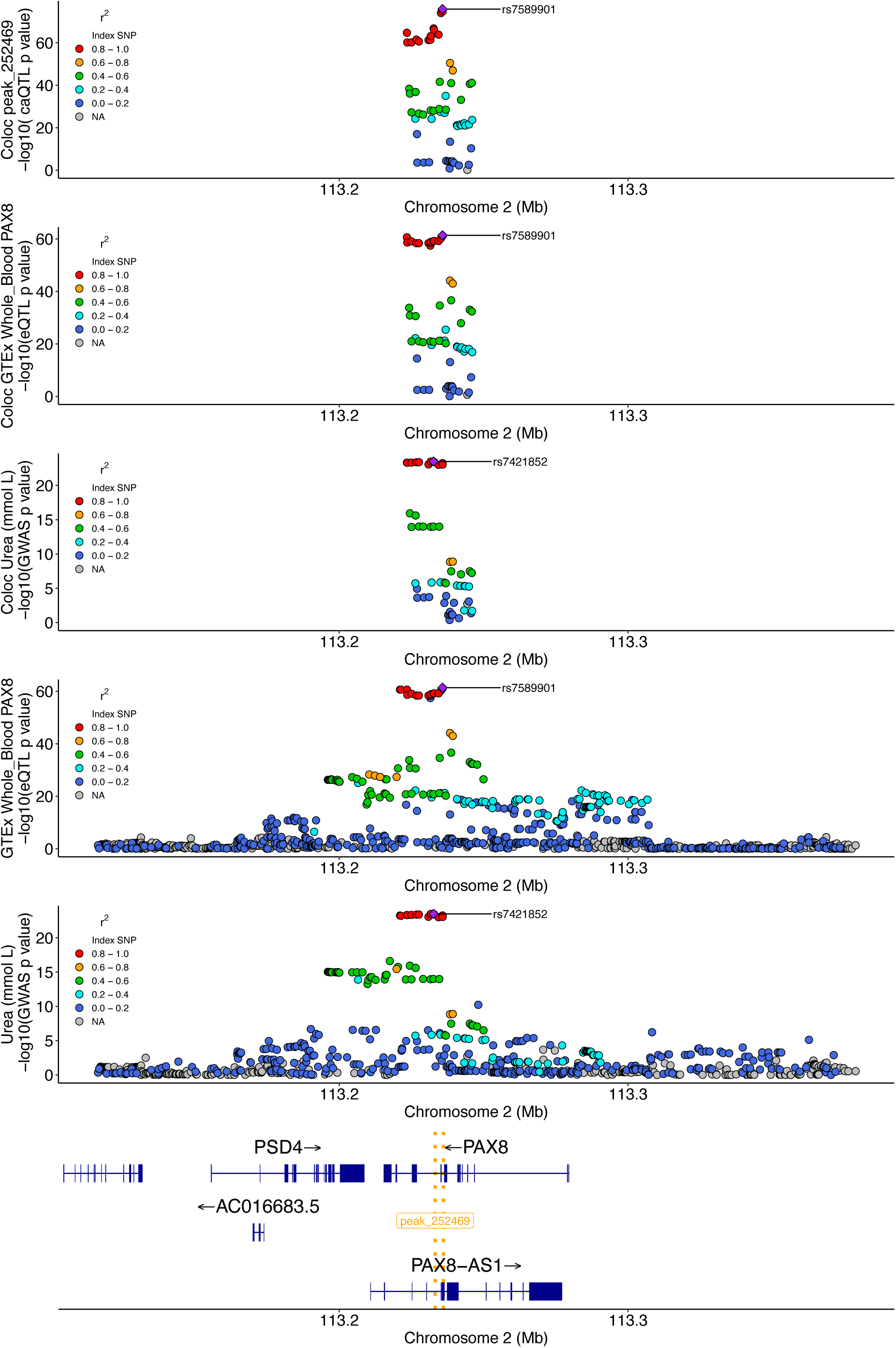
Change in chromatin accessibility and expression implicate *PAX8* in serum urea levels. The top three plots are the colocalization windows (10kb + caQTL peak) for the caQTL, eQTL, and GWAS, respectively. The following two plots are showing a larger window to illustrate the eQTL and GWAS signals, respectively, at this locus at a different scale. The bottom gene track highlights the position of genes at this locus, as well as the location of the caQTL peak (gold dotted lines).

### Sample heterogeneity enables identification of context-specific clusters

Because profiles of chromatin accessibility often segregate context or cell-type specific information, we next grouped our samples by their profiles of chromatin[70]. We performed dimensionality reduction[71] and applied a semi-supervised clustering method[71] to identify groups of similar samples, identifying 11 clusters (Figure 6A). We used sample metadata to assign a label to each cluster, denoting biological origin. Overall, clustering appears to be mainly driven by the tissue or cell type from which the sample is derived (Supplementary Figures 16-17). For example, blood cell types appear to be grouped together or near each other in separate, but related clusters. In addition, we found other examples of clusters where nearly half of the samples are derived from a single tissue, such as pancreas. Annotating samples with other aspects of metadata, such as primary sample vs. cell line, or cancer vs. non-cancer samples, did not appear to explain clustering results (Supplementary Figure 18).

**Figure 6.**
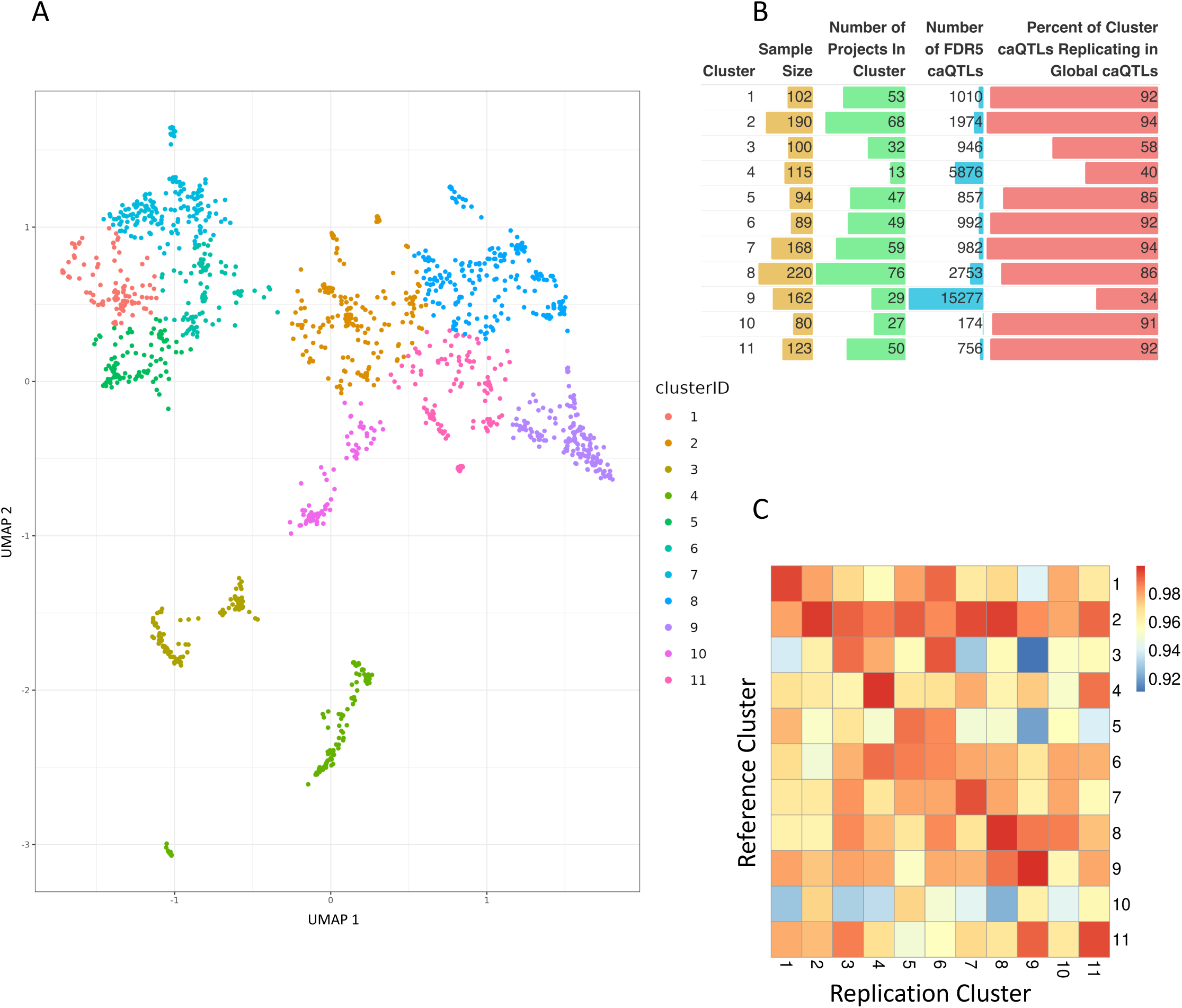
Clustering and discovery of cluster caQTLs across ATAC-Seq samples. **(A)** UMAP followed by k-means clustering to identify groups of related samples based on chromatin accessibility profiles across all peaks. **(B)** Cluster characteristics, caQTLs identified, and replication with respect to global caQTL mapping. **(C)** Replication rate (π_1_ value) of caQTLs identified in each cluster compared to those found in all other clusters.

### Clustering allows for identification of caQTLs in specific clusters

To determine whether clustering samples of similar biological origin enables the discovery of additional caQTL signals, we next performed caQTL mapping within each cluster. Each cluster is composed of a different number of samples, with varying contributions from cell types and projects, which is reflected in the number of caQTLs identified in each cluster. Cluster sample size ranged from 80-220 samples (Supplementary Table 17) and resulted in 174-15,277 (FDR<5%) caQTLs identified in a single cluster. As in the global analysis, cluster caQTLs showed similar patterns of genomic region annotation enrichments (Supplementary Figure 19) and lead caQTLs were centered within the open chromatin peak tested (Supplementary Figure 20). Across all clusters, cluster caQTLs rediscovered 34-94% of caQTL peaks observed in the global analysis (Figure 6B) with median global caQTL replication rate of 0.99 (π_1_ value) across all clusters (Supplementary Figure 21). Analysis comparing cluster caQTL peak discoveries to other clusters resulted in a range of caQTL peak rediscovery (Supplementary Figure 22) but high replication rate across clusters (π_1_ value 0.91-0.99) (Figure 6C, Supplementary Table 18). This suggests that clusters are capturing common global signals, but some clusters are better powered at identifying caQTLs that might be cell/tissue-specific. For example, cluster 9, which identified the largest number of cluster caQTLs, is comprised of more than 50% LCL samples, many of which are from a single study (Supplemental Table). Approximately 2/3 of the caQTL peaks identified in cluster 9 are not identified as caQTL peaks in the global analysis performed across all tissues/cell types, suggesting that cluster 9 may be better powered to discover caQTLs more prevalent in LCLs and related blood cell samples. As a measure of reproducibility across experiments, we found that Cluster 9 caQTL lead variants were enriched for evidence of caQTL peak causality in the original study[34] that the majority of cluster 9 samples originate from (Supplementary Figure 23). These results suggest that as with eQTLs, future work increasing the sample size to examine cell/tissue-specific caQTLs is likely to capture novel caQTLs that will be useful for elucidating molecular mechanisms underlying GWAS signals.

Mapping caQTLs in clusters highlights the increase in caQTL discovery power of aggregating all samples across experiments, particularly for caQTLs that might be found across cell types. In our global analysis we identified 23,381 caQTL peaks, with a maximum of 5,169 of those also identified in a single cluster caQTL mapping experiment. This suggests that by considering all samples, we achieve greater than a 4.5X increase in caQTL discovery power for global caQTLs. Across all clusters, we identify 8,610 (37% of global) caQTL peaks that were also found in the global analysis and 14,795 caQTL peaks that were not found in the global analysis.

### Cluster-specific caQTLs can explain additional gene regulation and GWAS signal causality

We next performed colocalization analysis between GTEx eQTLs and the caQTLs identified within each cluster to determine if cluster-specific caQTLs appear to be involved in gene regulation as well. As in the cluster caQTL analysis, we find that the number of colocalizations found per cluster was commensurate with the number of caQTLs identified in each cluster. We find a maximum of 13,688 unique eGenes colocalizing in a single cluster, and a total of 16,833 unique eGenes colocalize when considering all clusters (Supplementary Tables 19-20). Compared to the global analysis, which identified a total of 17,471 unique colocalizing eGenes, 14,017 of which also colocalized in the cluster analyses, suggesting that the majority of colocalizing eGenes are identified across both analyses. As in the cluster caQTL analyses, we find that colocalizing eGenes are often shared across clusters (Supplementary Figure 24). Considering all cluster colocalization events, 7,789 total eGenes were found to uniquely colocalize in a single cluster, with 5,532 (71%) of these in cluster 9. Overall, we find a variable number of cluster-specific caQTL/eQTL colocalizations per cluster, many of which are shared across clusters.

Our previous analyses assessed the benefit of utilizing global caQTLs in GWAS colocalizations compared to eQTLs. In this analysis, we considered eQTLs that were discovered in experiments performed in single tissues, experiments that are much more likely to identify variants with tissue-specific effects compared to our multi-tissue, global caQTL mapping strategy. Cluster-specific caQTLs might more closely mimic these single-tissue eQTL datasets, as these caQTLs were mapped in clusters of samples that likely shared a similar biological origin. To better compare the contribution of eQTLs and caQTLs to GWAS signals, we considered caQTLs identified in both global and cluster-specific analyses to assess colocalization improvement. Across all GWAS traits and eQTL tissues tested, we find that combining global and cluster-specific caQTLs results in an increase of the contribution of caQTLs to GWAS colocalizations. Specifically, we find a median of 41 GWAS signals colocalizing with caQTLs only and a median of 67.5 GWAS signals colocalizing with both caQTLs and eQTLs (Supplementary Figure 25, Supplementary Table 21). Both measurements are increases compared to the global analysis only. In contrast, the median number of GWAS signals that colocalize with eQTLs only decreased to 39 (Supplementary Figure 25, Supplementary Table 21). Leveraging both global and cluster caQTLs, together with eQTLs, we explained a median of 62% of GWAS signals tested (Supplementary Figure 26). Overall, we find that both global and cluster-specific caQTLs can contribute to the causal mechanisms underlying GWAS signals not captured by eQTLs.

## Discussion

We developed a pipeline to discover caQTLs on a large scale by aggregating and genotyping large-scale ATAC-seq data across many studies. We collected 10,293 human ATAC-seq samples, representing 1,454 unique donors, from public databases that come from a diversity of cell types and conditions, demonstrating that genotype data can be accurately called from ATAC-seq data, and identified unique sample donors, both within and across projects.

Combining accessibility and genotype information, we performed caQTL analysis and were able to capture global and cluster-specific caQTLs. caQTL studies are often limited by sample size constraints. We show that amassing public-domain project data allows for identification of a greater number of caQTLs than smaller individual studies alone. We demonstrated that caQTLs are enriched for various regulatory elements and likely underlie gene expression differences and complex human traits. We provide our large catalog of global and cluster caQTLs as a resource.

Our study does have limitations and opportunities for further development. Naturally, as more ATAC-seq data are generated, a similar study could be repeated on a larger scale.

Additionally, the clustering performed in our study was coarse, and may have grouped multiple cell types or contexts together. With a larger sample size from new studies or more extensive exploration of clustering methods or cell type prediction approaches, these grouping could be further refined and made more homogeneous, which would be expected to boost statistical power for discovery. Although we analyzed a large and diverse set of samples and experiments, many GWAS signals were not tagged by one of our caQTLs (and/or by eQTLs). One explanation for this is that we are missing many cluster/context-specific caQTLs that may underlie the remaining GWAS signals. One limitation of this study is that while the sample contexts were diverse, we still do not have sufficient sample size across some disease-relevant contexts to fully examine context-specific caQTLs. Further work, perhaps using single cell ATAC-seq data, is necessary to gain insight into tissue/cell context specific caQTLs. Other types of molecular QTLs may underlie some unexplained GWAS signals[60]. Incorporating additional data modalities, such as those reflecting chromosome conformation changes, may identify additional QTLs underlying GWAS loci. A recent study has shown that genetic variants in enhancer regions affect gene expression changes via enhancer-promoter touching and looping processes[72]. Integrating HiC or HiChIP datasets with ATAC-seq data can provide insight into this process. These datasets may also help identify target genes or resolve situations where multiple eGenes are implicated as causal genes at a locus[73]. Furthermore, other mechanisms, such as DNA methylation (meQTLs)[74,75] or post-transcriptional processes such as splicing (sQTLs)[66] or protein concentrations (pQTLs)[76] could underlie GWAS signals that have yet to be explained.

Although we observed colocalization analysis between our caQTLs and GWAS signals on par with previous studies [77], experimental validation is necessary to determine whether putative causal variants underlying these QTLs directly mediate disease risk[78,79]. Previous studies have shown that this type of analysis has led to the correct identification of molecular mechanisms underlying disease. For example, regulatory mapping has successfully identified gene targets that can be experimentally modulated to produce a phenotypic effect both in vitro and in vivo[80]. Furthermore, caQTL analyses have been used to predict mechanisms underlying GWAS signals with follow-up functional experiment results supporting these predictions[15]. Ultimately, regulatory elements and gene targets that we identify as implicated at GWAS loci will need additional support from low-throughput experimental techniques to confirm our findings, such as using base editing to dissect variant function[81]. Toward the goal of understanding molecular mechanisms underlying GWAS signals, molecular QTLs generate hypotheses and our work has demonstrated that including caQTLs in these experiments increases the number of GWAS signals for which a putative molecular mechanisms may be identified.

### Conclusions

In summary, we have deployed a pipeline to call a set of consensus peaks from thousands of publicly available ATAC-seq samples and genotype these samples directly from the experimental sequencing reads. We leveraged these data to identify caQTLs that likely share causal variants with eQTLs and GWAS signals. We show that caQTLs can improve our understanding of the mechanisms underlying GWAS signals and we provide this dataset as a resource for use in further fine-mapping experiments.

## METHODS

### Sample Collection

ATAC-seq samples were identified through the Gene Expression Omnibus (GEO) database and downloaded. Collected sample metadata is found in Supplementary Table 1.

### Benchmarking on HapMap samples

We downloaded ATAC-seq for 71 HapMap samples from ENA project PRJEB28318[34]. We aligned the sequencing reads to GRCh38 using bowtie2 and retained only autosomal chromosomes. Duplicated reads tagged by Picard were removed and Base Quality Score Recalibration (BQSR) was performed using GATK tools. Variant calling was done using GATK HaplotypeCaller. Loci with less than 2 reads were filtered out and variants were mapped to GRCh37 using Picard LiftoverVcf. Minimac4 was utilized to run imputation with reference panel derived 1000G Phase 3 (https://csg.sph.umich.edu/abecasis/mach/download/1000G.Phase3.v5.html). We kept only the genotype for common variants derived from 1000G with MAF > 0.05. The gold standard variants were obtained from https://www.internationalgenome.org/data-portal/data-collection/grch38. For the ATAC-seq data, we converted cram files to bam files, and removed the reads that map to mitochondrial genome. We obtained the genotype from the 1000 Genome Project on the GRCh38 genome assembly[82].

### Benchmarking for caQTLs in HapMap samples

We first obtained caQTLs using ATAC-seq reads with BH corrected P-value < 0.05, then ran QTL analysis using gold standard genotype and obtained caQTLs with BH corrected P-value < 0.05. The precision is computed as the percentage of replicated caQTLs at FDR < 0.05 using the gold standard genotype. Similarly, we first obtained caQTLs using gold standard genotypes with BH corrected P-value < 0.05, then ran QTL analysis using ATAC-seq reads and obtained caQTLs with BH corrected P-value < 0.05. The recall is computed as the percentage of replicated caQTLs at FDR < 0.05 using the ATAC-seq reads.

### Variant calling

For the ATAC-seq data, we performed two pipelines of variant calling, one using GATK HaplotypeCaller, and the other with Gencove’s low-pass sequencing pipeline. Using the GATK HaplotypeCaller, we performed alignment using Bowtie2, and removed duplicated reads and applied base quality score recalibration, followed by GATK HaplotypeCaller[33,83,84]. Variants with at least 3 reads were extracted. We then compared the called genotype dosage to the gold standard genotype by computing the Spearman correlation and mean squared error (MSE).

### Peak Calling

Genrich[27] (v0.6.1) was used to call peaks. A slightly modified version of Genrich was applied to allow peak calling across a large number of samples (https://github.com/maxdudek/Genrich). Genrich assigns p-values to genomic positions within each sample followed by combining p-values across samples using Fisher’s method to call peaks. Bam files were filtered using:

‘samtools view -S -b -q 10’. Bam files were name sorted using: ‘samtools sort -n

/path/to/q10_filtered_bams/sample.bam | samtools view -h -o

/path/to/nameSortedBams/sample.bam’. Peak calling parameters were: ‘Genrich -t

/path/to/nameSortedBams/sample1.bam,

path/to/nameSortedBams/sample2.bam, path/to/nameSortedBams/sampleN.bam, -j, -o /path/to/outputFile -v -E

/path/to/blacklistRegions.bed -r -q 0.05’.

### Genomic Annotation Enrichment

Genomic annotation enrichment analyses were performed using the R package annotatr (v.1.28.0) (https://bioconductor.org/packages/release/bioc/html/annotatr.html). 100 iterations of random, matched background data using bedtools shuffle with flags “-chrom -excl /path/to/blacklistRegions.bed -g /path/to/chrSizes.txt”. P values were calculated by quantifying the number of random data iterations that were more extreme than the true data values for each category.

### Encode Roadmap Enrichment

Histone ChIP-seq data derived from adult human samples were downloaded from https://www.encodeproject.org/search/?type=Experiment&status=released&award.project=Roadmap. ATAC-seq peaks that overlapped histone mark data were identified using bedtools intersect -wo -a /path/to/encodeData.bed -b /path/to/peakCoords.txt. 100 iterations of random, matched background data using bedtools shuffle with flags “-chrom -excl /path/to/blacklistRegions.bed -g /path/to/chrSizes.txt”. P values were calculated by quantifying the number of random data iterations that were more extreme than the true data values for each histone mark.

### caQTL Mapping

Sample peak counts were generated for all samples. To remove potential outlier peak regions, peaks with mean count <1 and max count> 100,000 were removed. Peaks were also removed if >5000 samples had a read count of zero in that peak. Given that a single individual might contribute multiple samples to the 10,293 sample pool, we identified each sample that can be attributed to each individual and averaged sample peak CPM values to calculate a single CPM value per peak for each individual donor. This workflow results in 1454 individual donor samples for caQTL mapping. Code available in file “Post_peakCalling_CountMatrixGeneration_Pipeline.txt”. tensorQTL (v.1.0.9) [85] was used to identify caQTLs using a linear model with 3 genotype PCs and 200 principal components as covariates. PCs generated from each cluster’s chromatin accessibility peak read count data

sample matrix was used to map caQTLs on chromosome 1 over a large range of included PCs. The optimized PC covariate number was chosen based on the elbow of the PCs included vs. caQTL discovery plot on chromosome 1 (Supplementary Table 23). We tested all genotyped biallelic genetic variants with MAF > 0.05 within 10 kilobases of all open chromatin peak boundaries detected by Genrich from the ATAC-Seq data[35]. Empirical p-values were estimated by tensorQTL to get peak-level p-values and q-values [86]. caQTL mapping code available in file “caQTL_mapping_code_pipeline.txt”.

### Lead caQTL/eQTL Enrichment

Significant lead eQTL variants were downloaded for 49 tissues from GTEx v8 publicly available data. Unique global sample analysis lead caQTLs (n= 21,647) were intersected with lead eQTL variants to assess overlap within each GTEx tissue. The unique intersection of overlaps across all tissues was considered to determine the total number of caQTL lead variants that were found to be a lead eQTL variant in at least one tissue. Background variants were selected to perform enrichment analyses. Background variants were chosen by randomly sampling non-lead caQTL genetic variants that were matched, +/- 10%, to the allele frequency and distance to nearest gene transcription start site of true lead caQTL variants. Enrichment of caQTLs/eQTLs in each tissue was calculated as the ratio of the overlap of true lead caQTL/eQTL compared to the overlap of background variants/eQTL across 100 iterations.

### Replication Analysis

An external dataset was identified that was not included in our peak calling or caQTL mapping workflow[49]. Global FDR5 caQTL peaks with any overlap with the external study and variants tested in both analyses against these shared peaks were identified. External study p values were used for π_1_ replication rate calculation and plotted.

### GWAS Trait/Signal Selection

GWAS summary stats for traits were downloaded February 2021 from the UKBB Neale Lab repository and selected for relevant traits based on the following filters: h2 > 0.05, z > 7, confidence == high. Independent significant GWAS signals from 78 traits were chosen to prevent counting a single GWAS signal multiple times. This was done by selecting GWAS signals with a minimum p-value of 5e-08, considering a window of 50 kb on either side of these variants, clumping all variants with R2 > 0.01, and selecting the variant with the most significant p-value as the lead GWAS signal for this locus.

### Colocalization Analyses

Colocalization was performed using coloc[59] (v.5.2.3). All reported colocalizations utilized a previously published approach to define significance[87]. This approach consists of considering whether the colocalization is sufficiently powered, PP3+PP4 > 0.8. For those events that surpass this threshold, we assessed whether the colocalization is significant, PP4/(PP3+PP4) > 0.9. GTEx v8 data were downloaded from https://www.gtexportal.org/home/downloads/adult-gtex/bulk_tissue_expression.

### Colocalization Genome Annotations

Genomic annotation enrichment analyses were performed using the R package annotatr (v.1.28.0)(https://bioconductor.org/packages/release/bioc/html/annotatr.html). For each type of colocalization, caQTL peaks involved in the colocalization were labeled with genomic annotations they overlap. To perform an enrichment analysis, true data results were compared with the median of 1000 iterations of random genomic regions matched to the true data using bedtools shuffle with flags “-chrom -excl /path/to/blacklistRegions.bed -g /path/to/chrSizes.txt”. Summaries were produced by identifying significant enrichments (annotation category enriched/depleted p value <= 0.05) across all traits or trait/tissue pairs and calculating the mean and median enrichment/depletion values.

### Clustering Analyses

To reduce the dimensions of the data, Uniform Manifold Approximation and Projection (UMAP) was performed on the normalized sample CPM count matrix across all peaks. Kmeans clustering was performed on UMAP coordinates 1 and 2. 11 outlier samples were removed from analysis. The number of clusters was optimized using several clustering metrics (Supplementary Table 22) and samples were assigned to a cluster based on the results of the clustering algorithm.

### Cluster-specific caQTL mapping

caQTL mapping was performed as in the global analysis. In this analysis, peaks identified in the global analysis were included if at least 50% of cluster samples had non-zero CPMs in that feature, resulting in the removal of 5-5920 (0.0003-0.35% of total peaks). All steps of the caQTL mapping pipeline were performed within each cluster. caQTL mapping was performed including 3 genotype PCs and an optimized number of principal components based on each cluster. For each cluster, a range of PCs generated from each cluster’s chromatin accessibility peak read count data sample matrix was used to map caQTLs on chromosome 1. The optimized PC covariate number was chosen based on the elbow of the PCs included vs. caQTL discovery plot. We tested all genotyped biallelic genetic variants with MAF > 0.05 within 10 kilobases of all open chromatin peak boundaries detected by Genrich from the ATAC-Seq data[35]. Empirical p-values were estimated by tensorQTL to get peak-level p-values and q-values [86]. All colocalizations were performed as described for the global analyses.

### Cluster caQTL replication analyses

Cluster caQTL replication of global caQTLs was assessed by extracting global caQTL peak test statistics from each cluster and calculating π_1_ replication rate. The reported replication rate for each cluster was calculated by calculating the median π_1_ replication rate after calculating π_1_ replication rate with a range of values for the lambda parameter (from=0.1,to=0.9,by=0.05).

Cluster caQTL replication rate across all other clusters was calculated in a similar fashion. For each cluster, cluster caQTL peak test statistics were extract from all other clusters and π_1_ replication rate was calculated. The reported replication rate for each cluster was calculated by calculating the median π_1_ replication rate after calculating π_1_ replication rate with a range of values for the lambda parameter (from=0.1,to=0.9,by=0.05).

## Declarations

### Ethics approval and consent to participate

‘Not applicable’

### Consent for publication

‘Not applicable’

### Availability of data and materials

All data generated or analyzed during this study are included in this published article [and its supplementary information files]. Publicly available samples used are listed in Supplementary Table 1. The code used to generate the results and figures and generated data/results are deposited in a Zenodo repository (https://doi.org/10.5281/zenodo.12706263) and will be made public upon publication.

## Authors’ contributions

A.B., C.D.B., Y.H., B.M.W. designed the study. J.K.P. and J.H.L. generated genotype data. Y.H. and B.M.W. performed computational analyses and prepared figures and tables. N.C. and T.L. assisted with analyses. M.F.D. assisted with software. A.B., R.K., B.F.V., Y.H. and B.M.W. wrote and revised the manuscript. All authors read and approved the final manuscript.

## Acknowledgements

‘Not applicable’

