## Supplementary Figures for "Genotype inference from aggregated chromatin accessibility data reveals genetic regulatory mechanisms": Supplementary_Figures_7.9.24.pdf

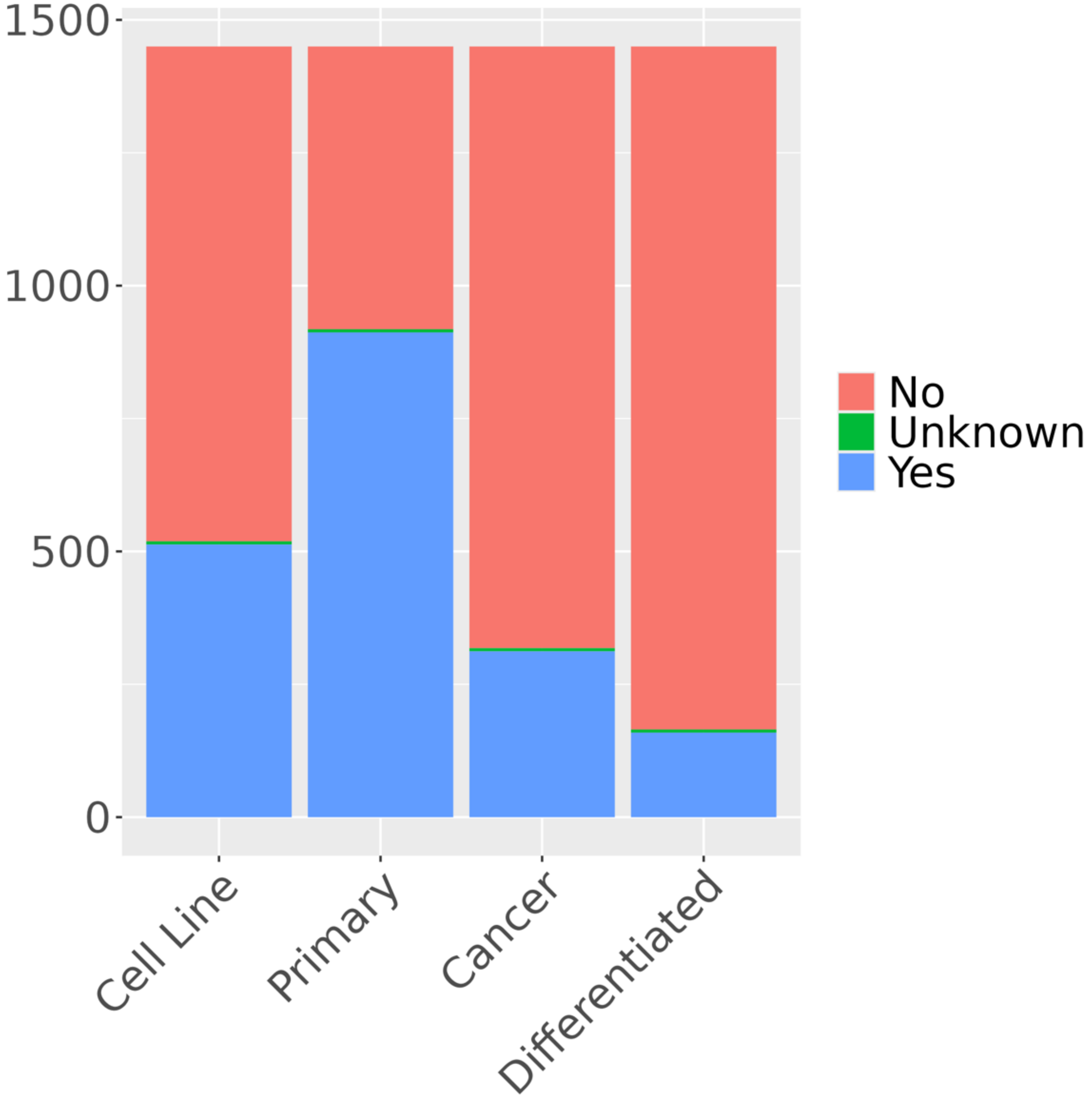


Supplementary Figure 1: Proportion of samples used for caQTL mapping, based on metadata review, from cell lines, primary tissue, cancer, and experimentally differentiated cell types.

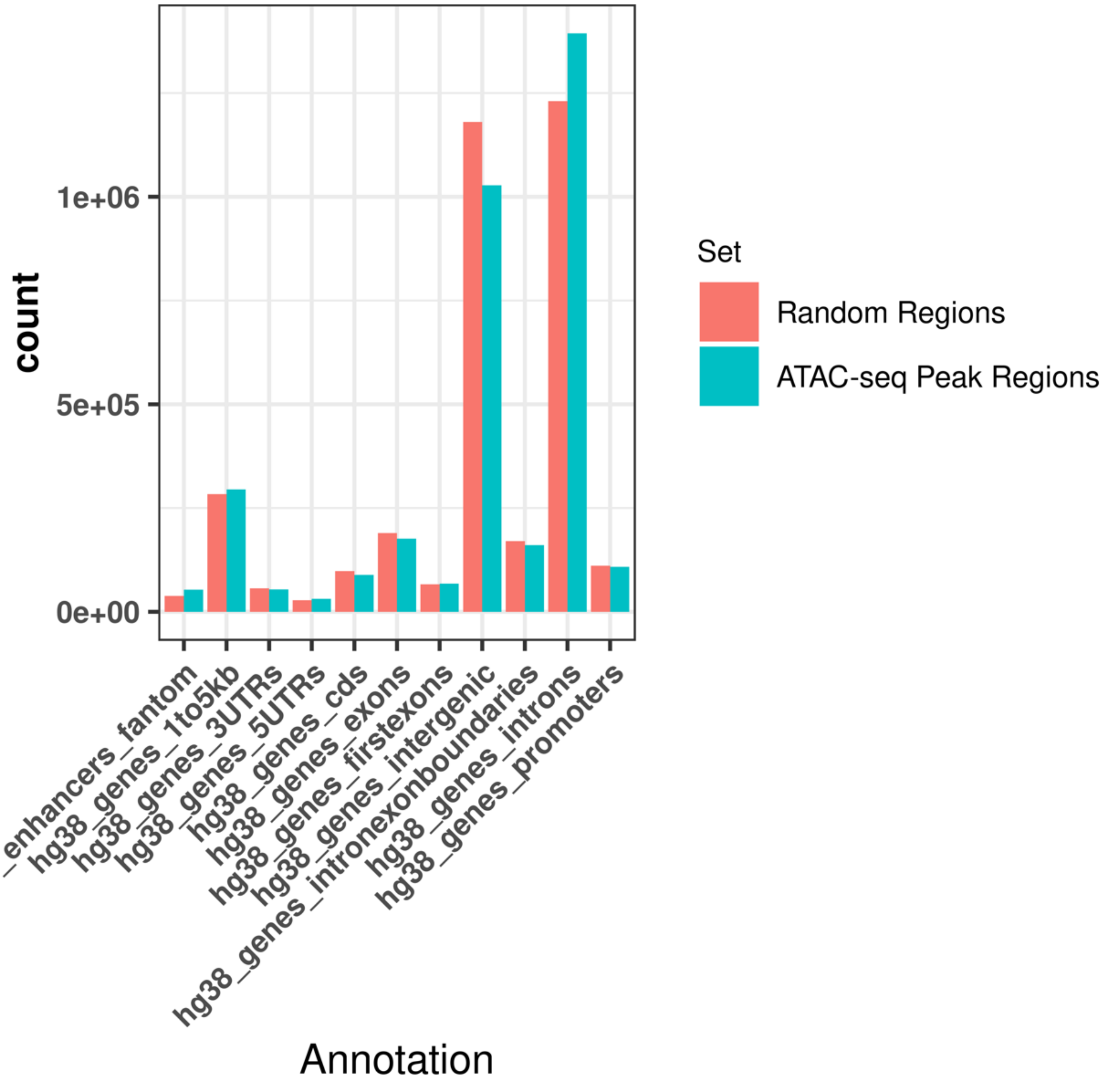


Supplementary Figure 2: Enrichment of all called peaks, compared to matched random control regions, in various genomic annotation categories.

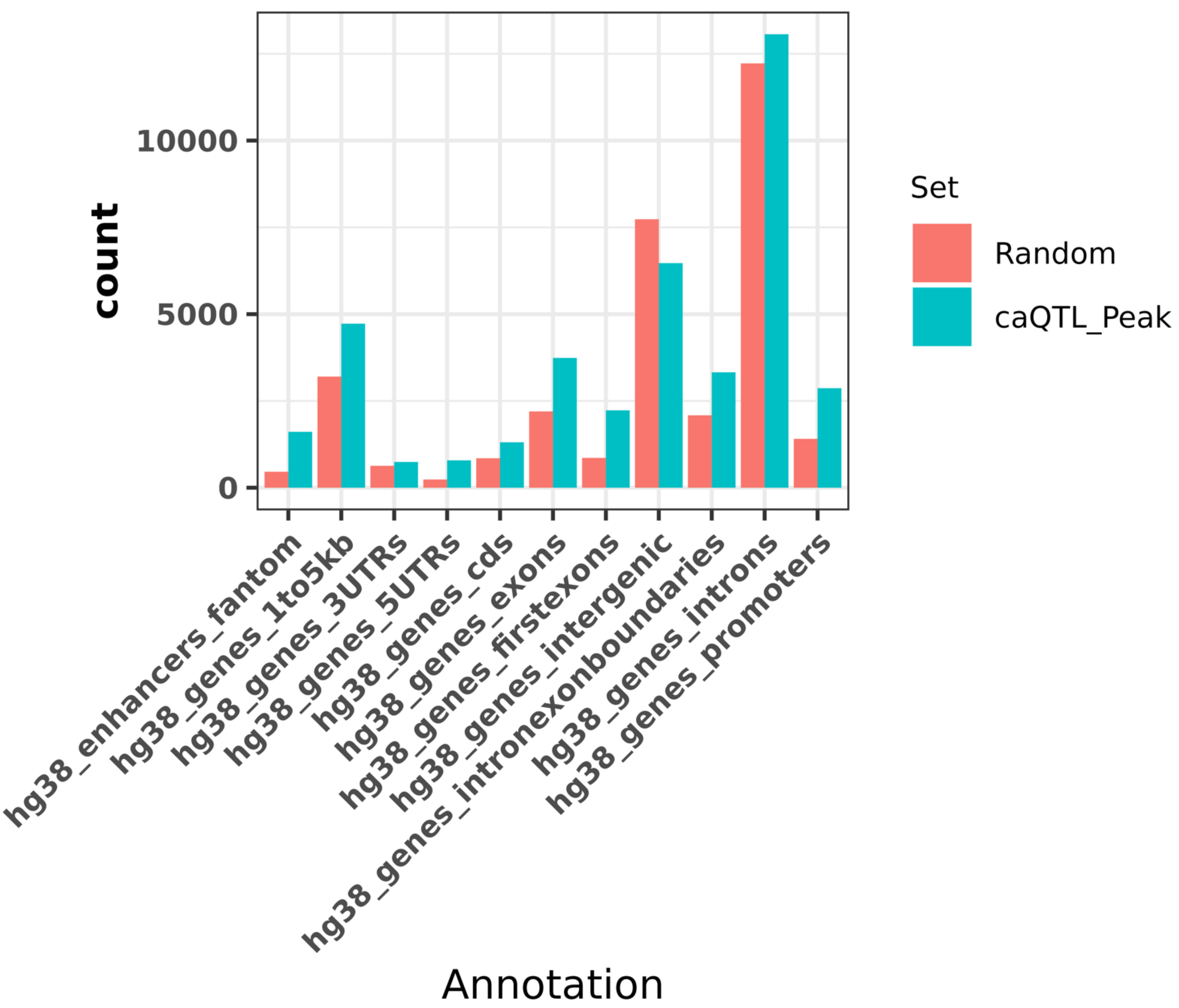


Supplementary Figure 3: Enrichment of caQTL peaks, compared to matched random control regions, in various genomic annotation categories.

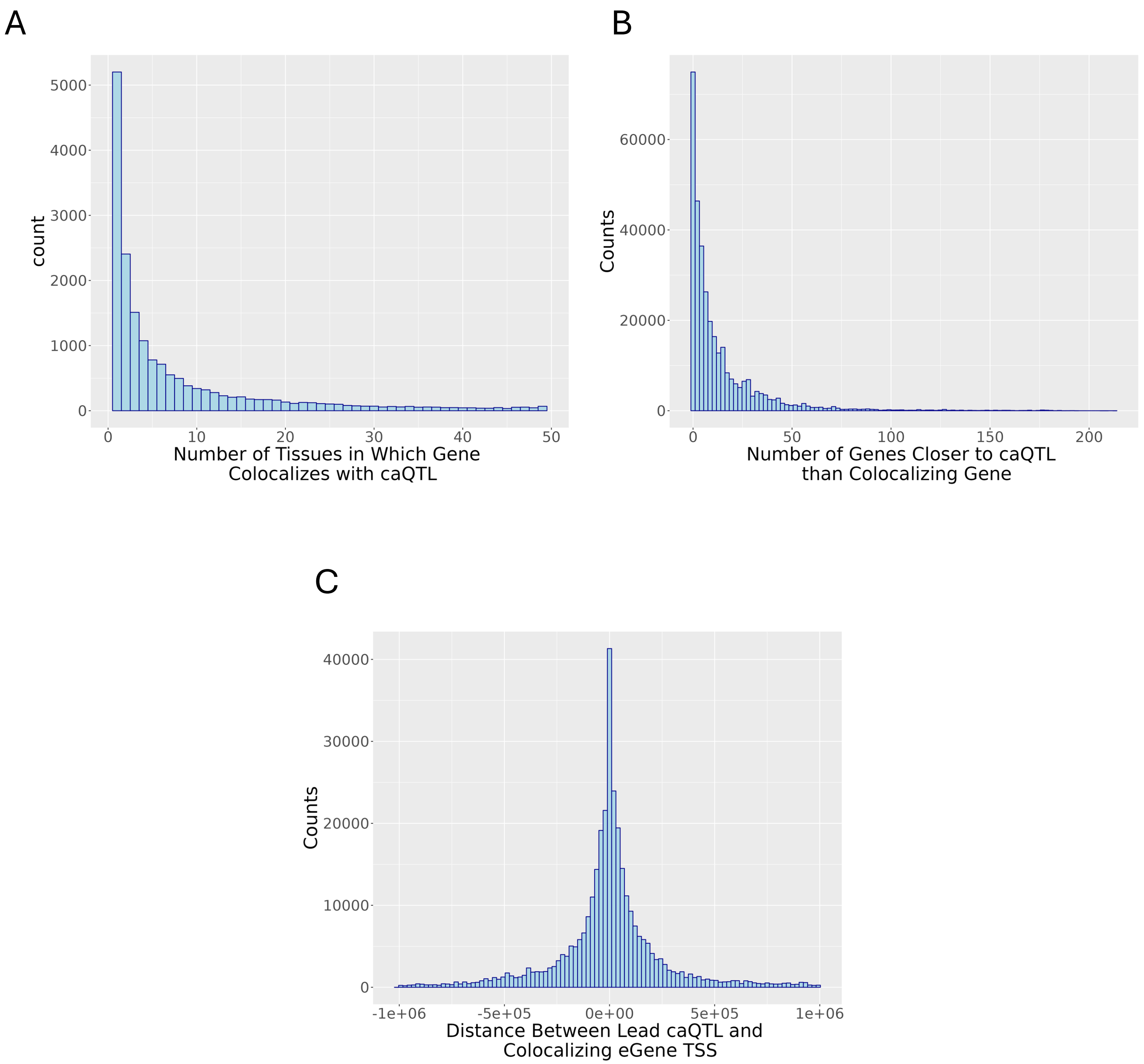


Supplementary Figure 4: A. Number of GTEx tissues in which gene colocalized with caQTL. B. Number of gene TSSs closer to lead caQTL than gene caQTL colocalizes with. C. Distance between the lead caQTL position and the TSS of the colocalizing gene.

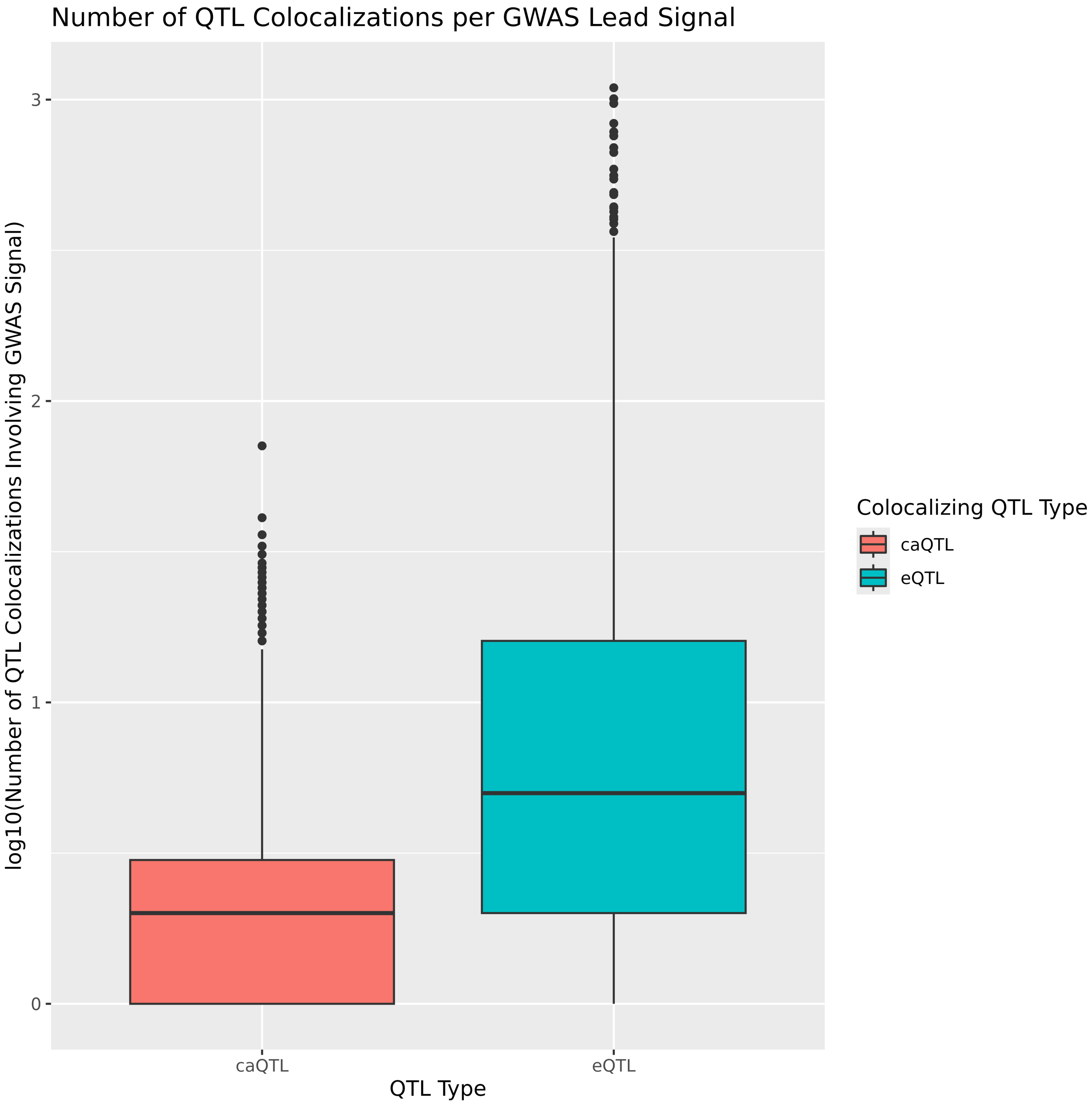


Supplementary Figure 5: Colocalizations were performed between caQTL/GWAS and eQTL/GWAS and number of filtered colocalizations for each GWAS signal plotted.

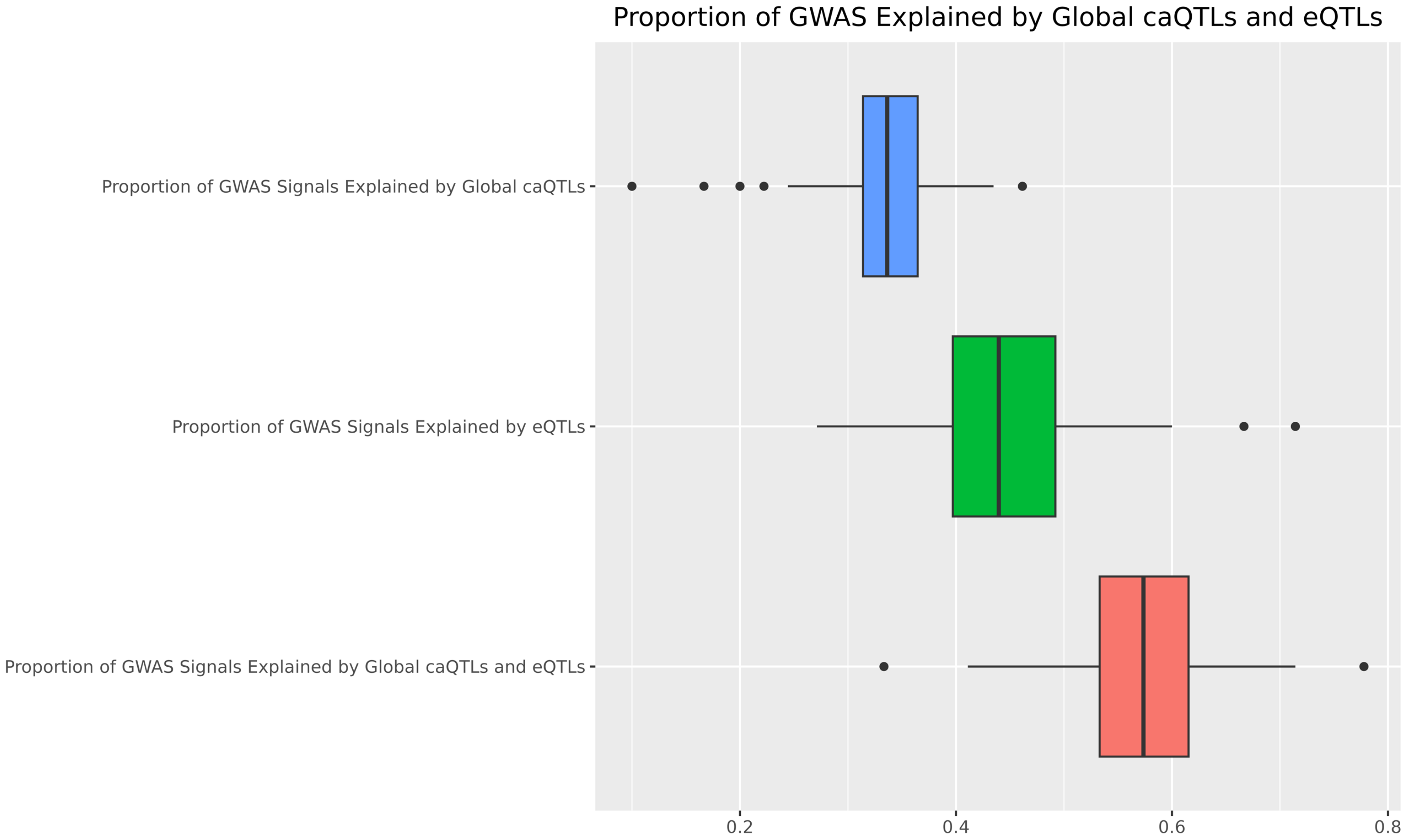


Supplementary Figure 6: Colocalizations were performed between caQTL/GWAS and eQTL/GWAS. Each GWAS signal was checked to see if it colocalized exclusively with caQTLs, eQTLs, or colocalized with both. Proportion of tested GWAS signals that colocalized in each category are plotted.

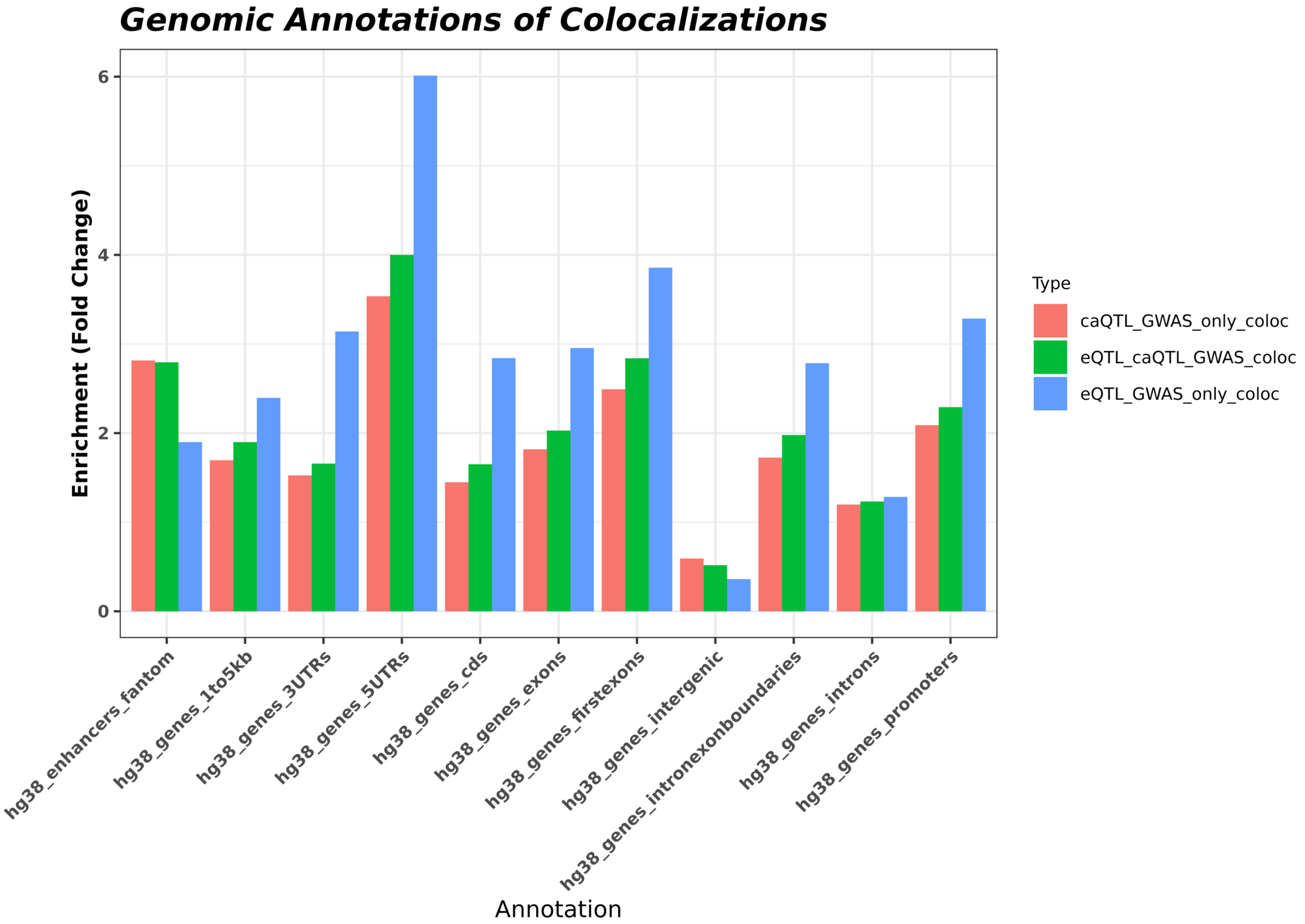


Supplementary Figure 7: Enrichment of all colocalization categories compared to matched random control regions, in various genomic annotation categories. Lead caQTL (+/- 250 bp) used for enrichments for colocalizations involving caQTLs and lead eQTL (+/- 250 bp) used for enrichments for colocalizations involving eQTLs.

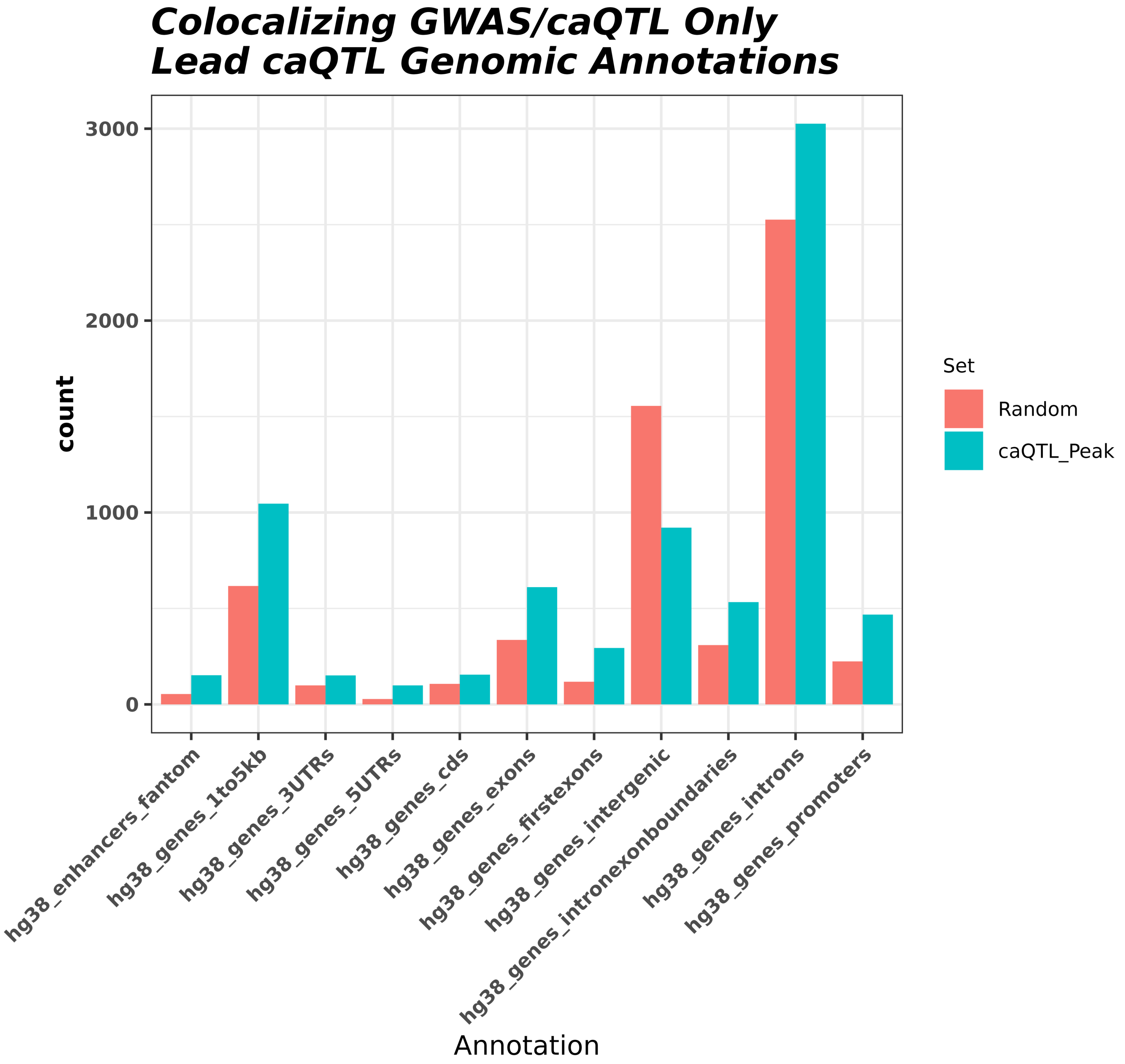


Supplementary Figure 8: Enrichment of lead caQTL (+/- 250bp) from caQTL/GWAS colocalizations only, compared to matched random control regions, in various genomic annotation categories.

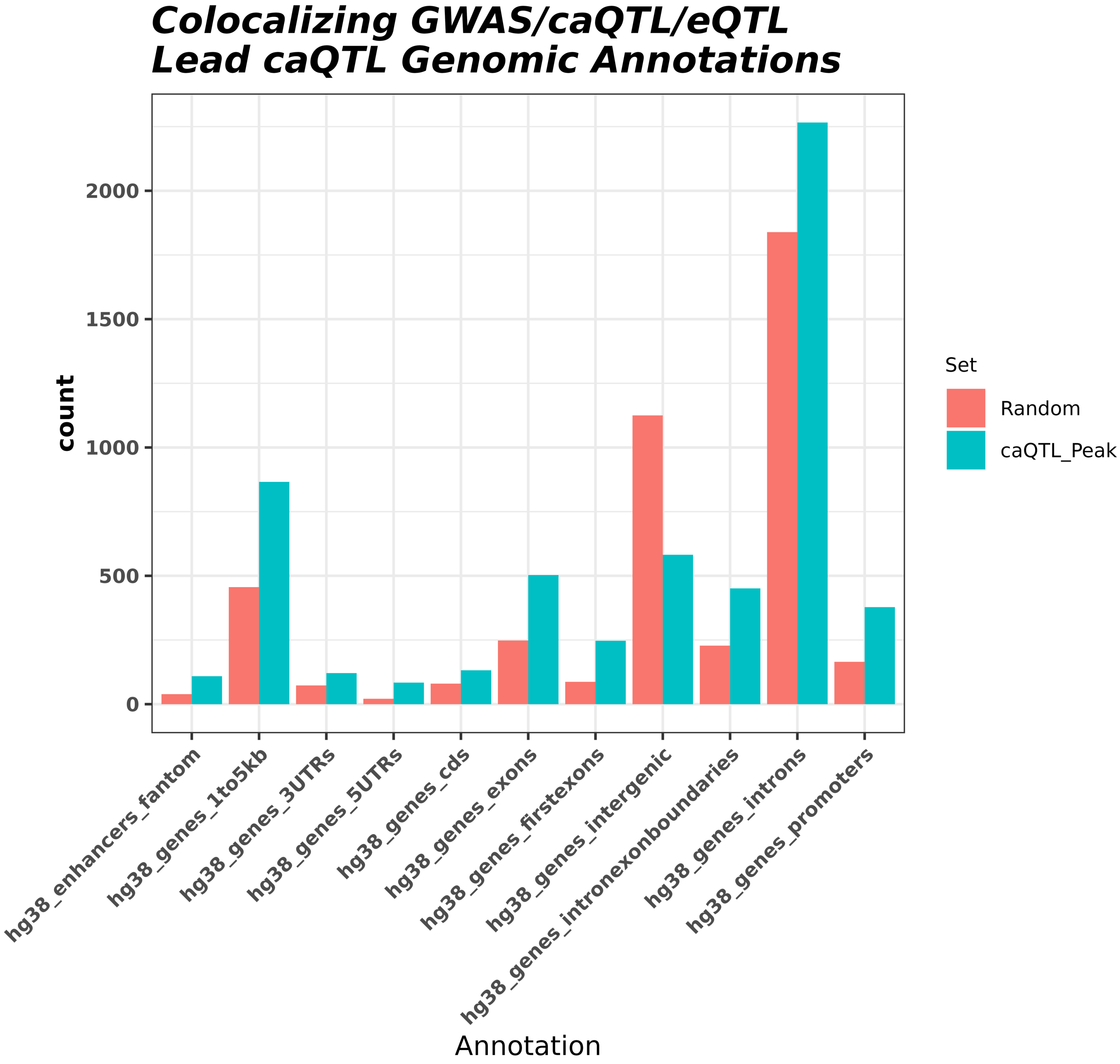


Supplementary Figure 9: Enrichment of lead caQTL (+/- 250 bp) from caQTL/eQTL/GWAS colocalizations only, compared to matched random control regions, in various genomic annotation categories.

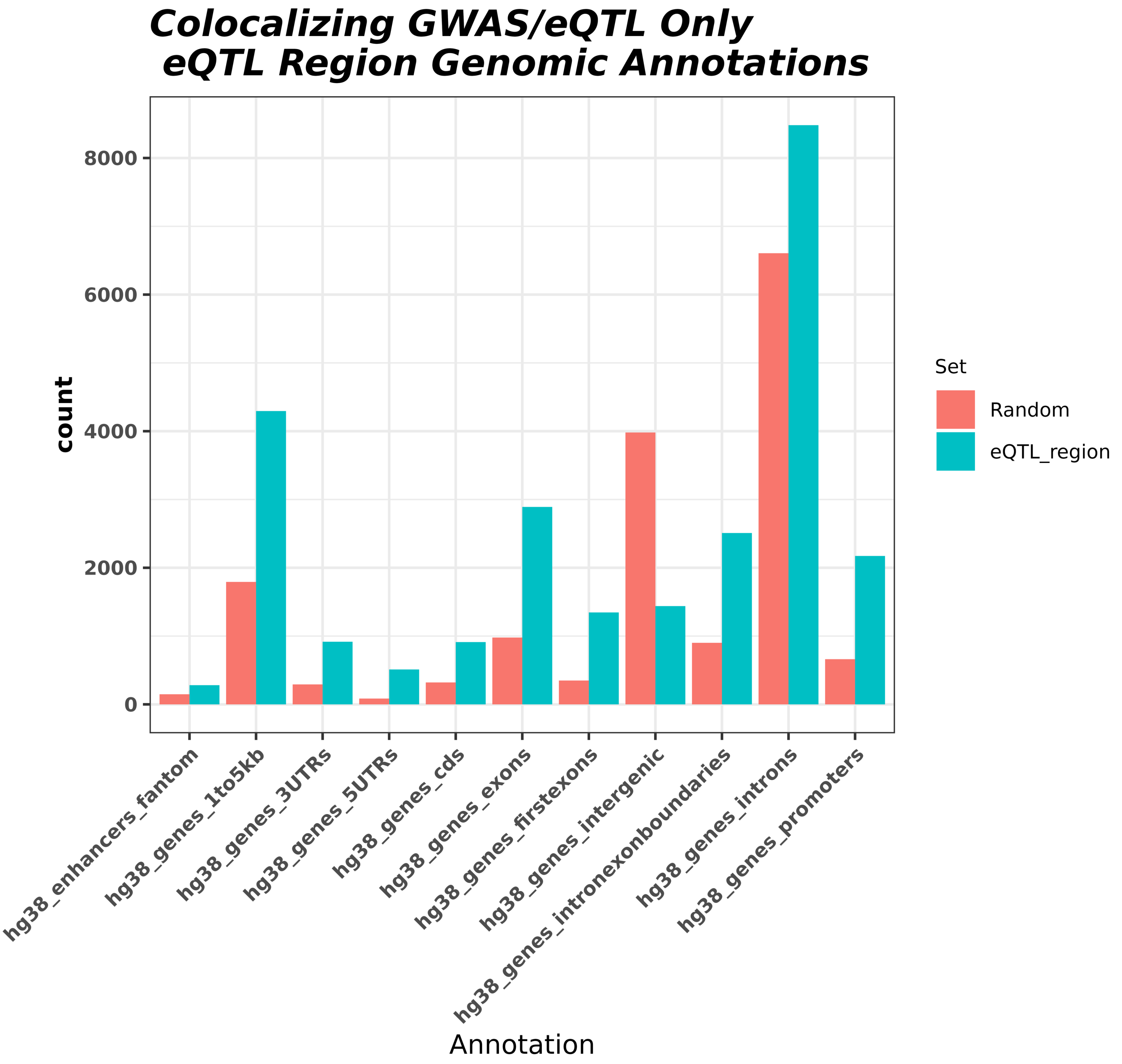


Supplementary Figure 10: Enrichment of eQTL (lead variant +/- 250bp) from eQTL/GWAS colocalizations only, compared to matched random control regions, in various genomic annotation categories.

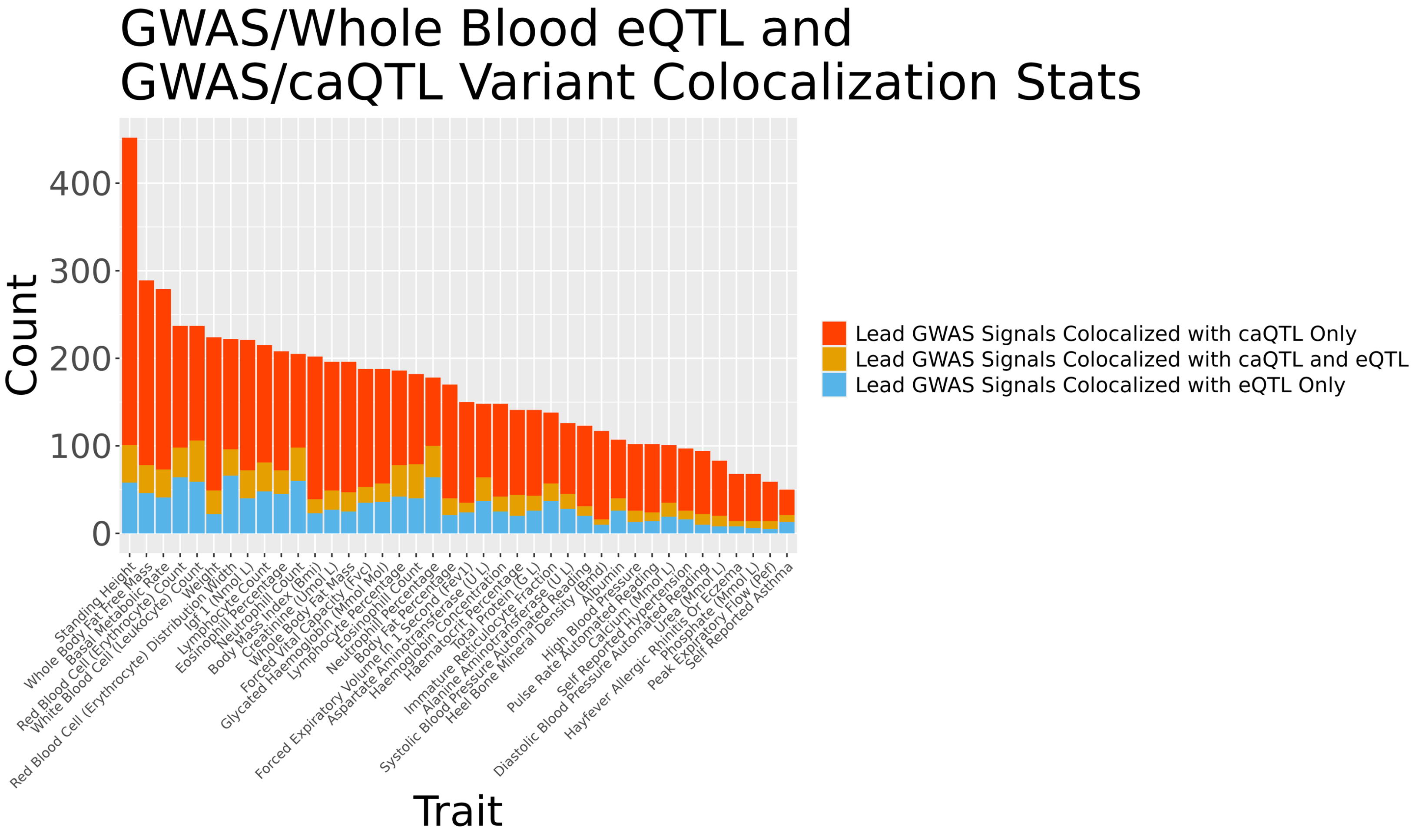


Supplementary Figure 11: For each GWAS trait, independent lead GWAS variant signals were checked for colocalization with caQTL and eQTL signals in GTEx Whole Blood only. Plotted is the number of unique lead GWAS variants per colocalization group, as multiple caQTL peaks, eGenes, etc. can colocalize with the same lead GWAS signal.

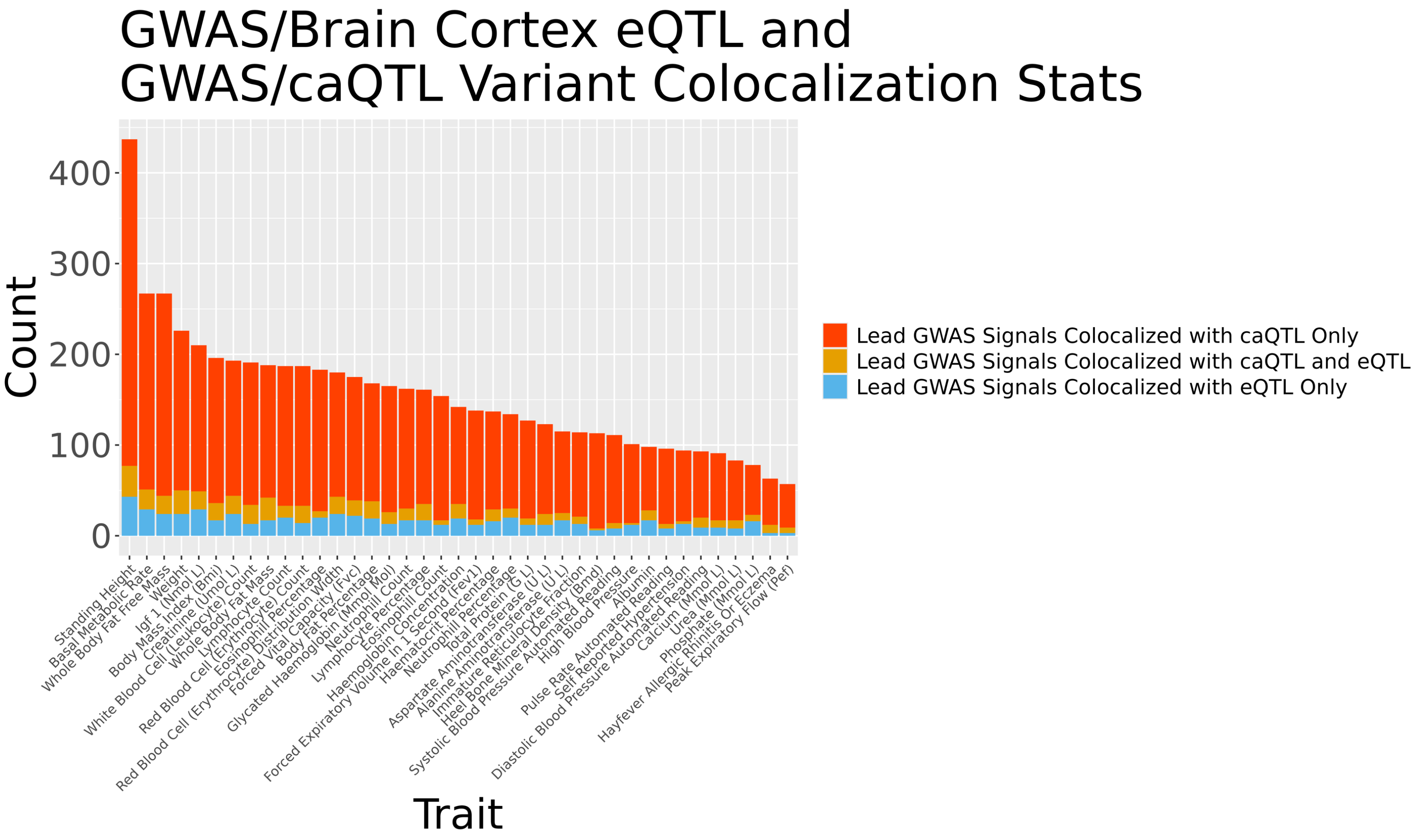


Supplementary Figure 12: For each GWAS trait, independent lead GWAS variant signals were checked for colocalization with caQTL and eQTL signals in GTEx Brain Cortex only. Plotted is the number of unique lead GWAS variants per colocalization group, as multiple caQTL peaks, eGenes, etc. can colocalize with the same lead GWAS signal.

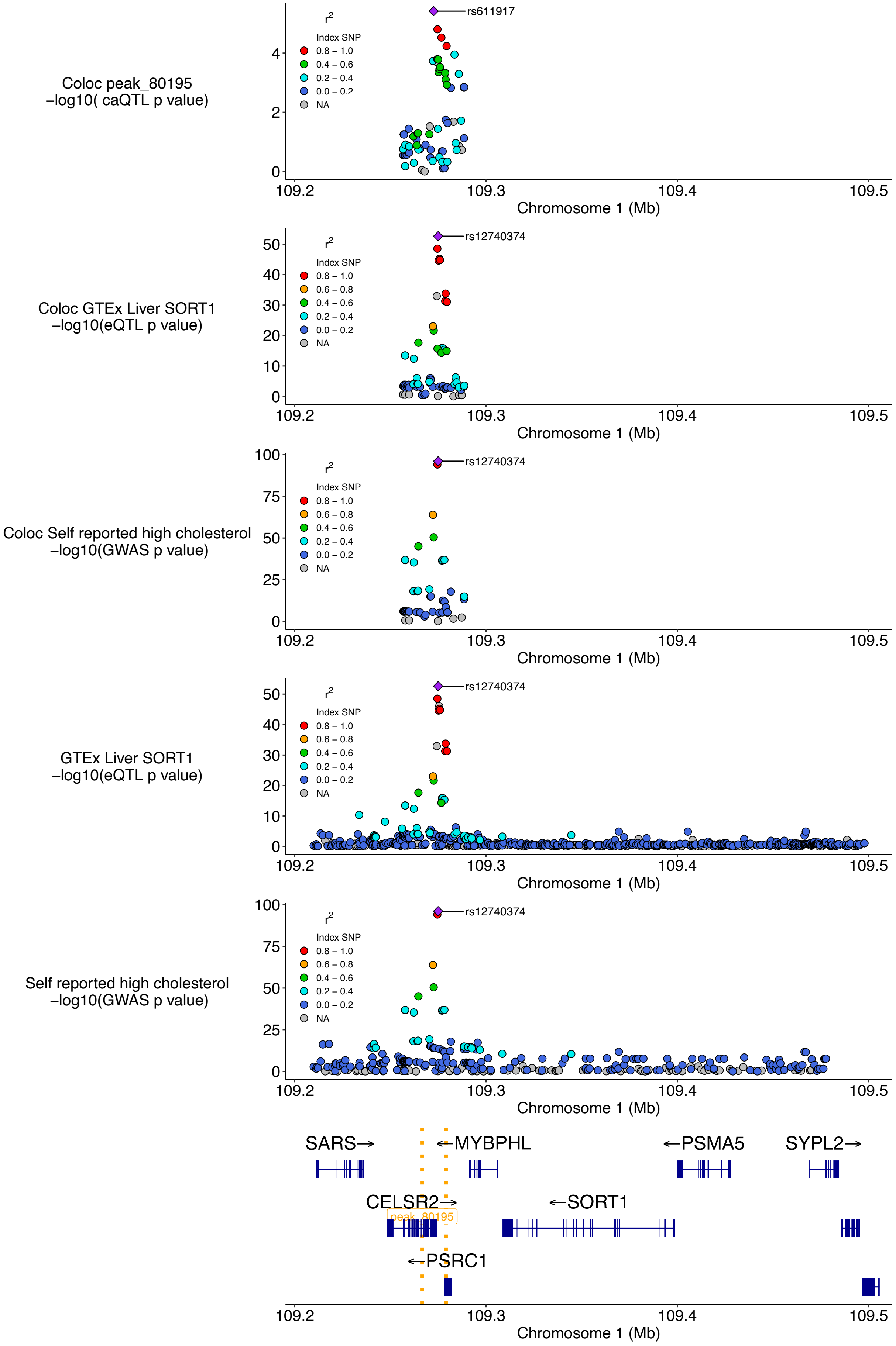


Supplementary Figure 13: caQTL peak at well-studied LDL GWAS locus colocalizes with *SORT1* in liver and GWAS signal for high cholesterol. The top three plots are the colocalization windows (10kb + caQTL peak) for the caQTL, eQTL, and GWAS, respectively. The following two plots are showing a larger window to illustrate the eQTL and GWAS signals, respectively, at this locus at a different scale. The bottom gene track highlights the position of genes at this locus, as well as the location of the caQTL peak (gold dotted lines).

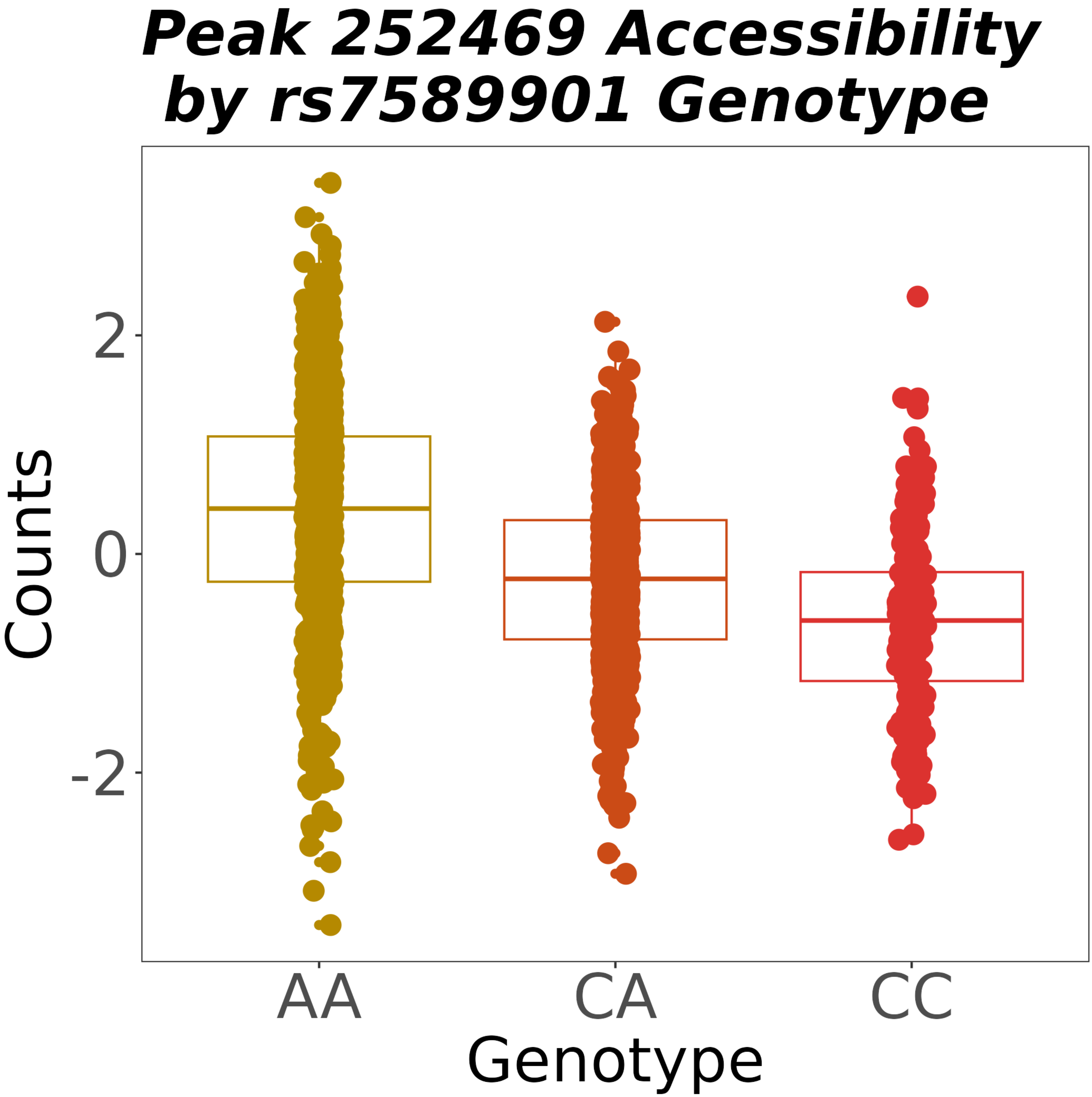


Supplementary Figure 14: Sample normalized read counts, grouped by genotype, at peak 252469, which colocalized with *PAX8* eQTL in Whole Blood and Urea serum levels. AA individuals have increased chromatin accessibility compared to CA and CC individuals.

### 3 models available for ZNF135 at chr2:113235767A>C.1

| Database | Model ID | WT score | MT score | WT start | WT end | MT start | MT end | WT strand | MT strand | Prediction | Score |
|---|---|---|---|---|---|---|---|---|---|---|---|
| jaspar2022 | MA1587.1 | 0.8607 | 0.9376 | -2 | 11 | -2 | 11 | plus | plus | gain | 0.3169 |
| jaspar2022DetailedTFFMs | TFFM0632.1 | 0.0309 | 0.8987 | -2 | 11 | -2 | 11 | plus | plus | gain | 0.9949 |
| jaspar2022FirstOrderTFFMs | TFFM0632.1 | 0.4564 | 0.9944 | -2 | 11 | -2 | 11 | plus | plus | gain | 0.9983 |

*(The combined score reflects only the TFFMs.)* **Combined** gain 0.9966

### Sequences

| | -14 | -13 | -12 | -11 | -10 | -9 | -8 | -7 | -6 | -5 | -4 | -3 | -2 | -1 | 0 | +1 | 2 | 3 | 4 | 5 | 6 | 7 | 8 | 9 | 10 | 11 | 12 | 13 | 14 | 15 | 16 |
|---|---|---|---|---|---|---|---|---|---|---|---|---|---|---|---|---|---|---|---|---|---|---|---|---|---|---|---|---|---|---|---|
| WT | G | G | G | G | A | G | A | G | A | G | A | A | G | C | T | A | G | A | C | C | T | C | C | C | T | G | C | T | G | C | G |
| MT | G | G | G | G | A | G | A | G | A | G | A | A | G | C | T | C | G | A | C | C | T | C | C | C | T | G | C | T | G | C | G |

### Known TFBSs at position

**ENCODE:** ATF7, BHLHE40, CBFB, CTCF, E4F1, ELF1, FOXK2, GMEB1, IKZF1, IKZF2, MAX, MGA, MNT, MTA3, MYC, NFIC, PML, RBFOX2, RELB, SIN3A, SMAD5, TAF1, YY1, ZFX

**Ensembl:** ATF7, E2F6, ELF1, ELF4, FOXK2, GMEB1, IKZF1, IKZF2, KLF5, MGA, MNT, MYC, NR2F1, POU2F2, RELB, SIN3A, SP1, TAF1, TBP, ZBTB7A

**FANTOM5:** -

### Sequence logos

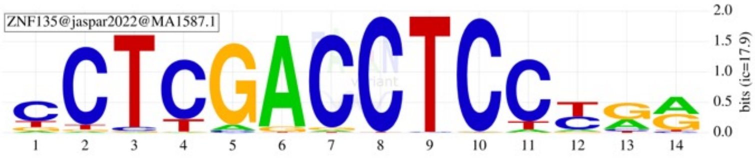


Motif display help

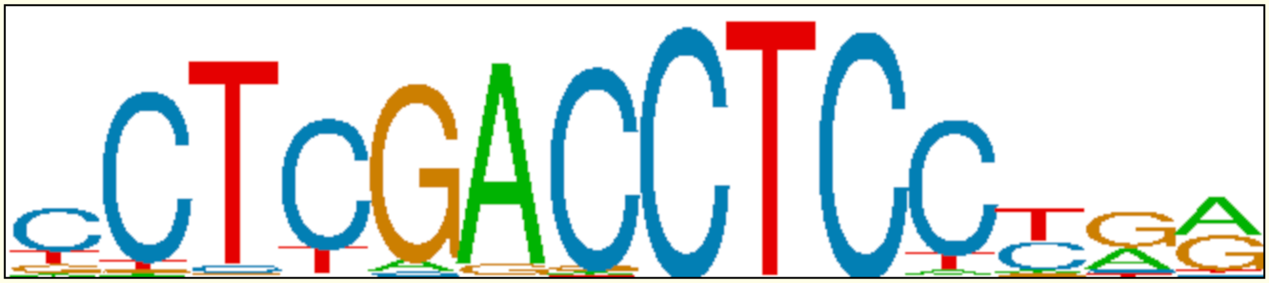

| | 1 | 2 | 3 | 4 | 5 | 6 | 7 | 8 | 9 | 10 | 11 | 12 | 13 | 14 | |
|---|---|---|---|---|---|---|---|---|---|---|---|---|---|---|---|
| | G | C | T | A | G | A | C | C | T | C | C | C | T | G | this occurrence |
| | c | **C** | **T** | C | **G** | **A** | **C** | **C** | **T** | **C** | C | t | g | a | motif consensus |
| A | 0.05 | 0.02 | 0.00 | 0.01 | 0.06 | 0.93 | 0.02 | 0.00 | 0.00 | 0.00 | 0.04 | 0.06 | 0.32 | 0.47 | |
| C | 0.63 | 0.91 | 0.04 | 0.81 | 0.03 | 0.00 | 0.95 | 0.99 | 0.01 | 0.99 | 0.84 | 0.42 | 0.07 | 0.04 | |
| G | 0.13 | 0.02 | 0.02 | 0.01 | 0.91 | 0.05 | 0.03 | 0.00 | 0.00 | 0.01 | 0.01 | 0.05 | 0.55 | 0.44 | |
| T | 0.19 | 0.05 | 0.94 | 0.17 | 0.00 | 0.01 | 0.01 | 0.01 | 0.99 | 0.00 | 0.11 | 0.47 | 0.06 | 0.05 | |

Supplementary Figure 15: Lead caQTL variant at caQTL/eQTL/GWAS colocalizing locus is rs7589901. Motif analysis predicts that ZNF135 will bind to motif overlapping rs7589901. The alternate ‘C’ allele at position 4 is predicted to increase binding affinity.

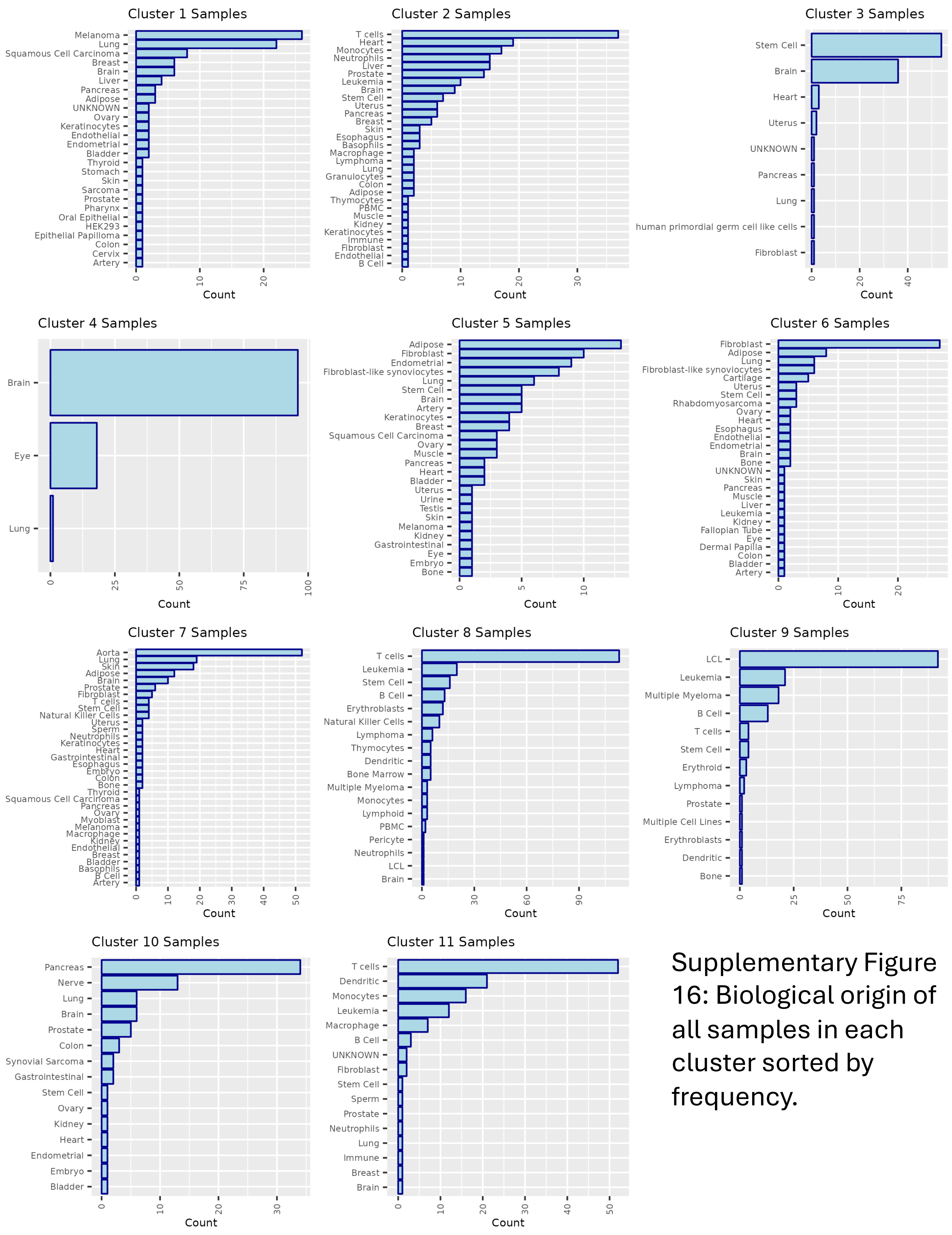


Supplementary Figure 16: Biological origin of all samples in each cluster sorted by frequency.

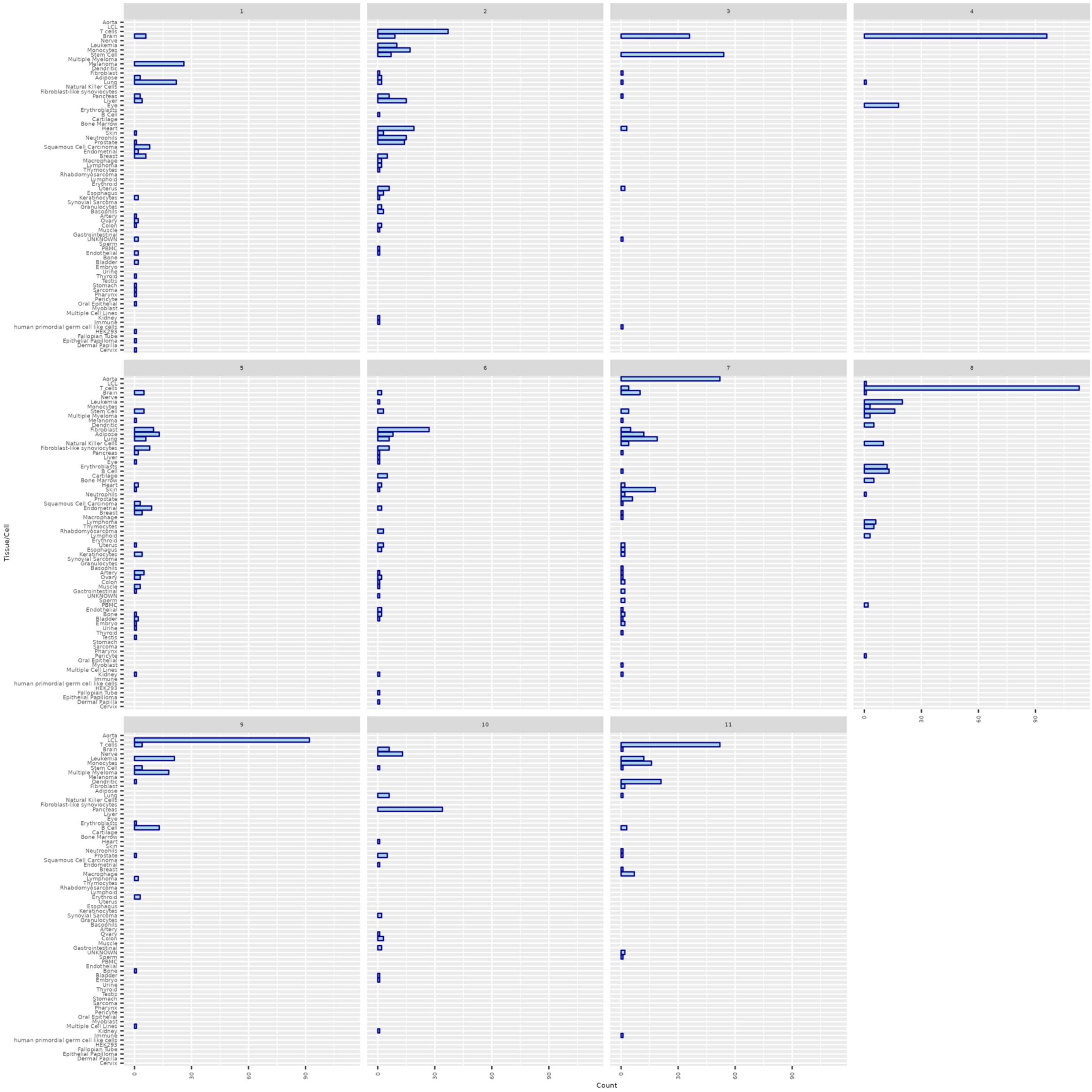


Supplementary Figure 17: Proportion of each cluster comprised of each biological sample type.

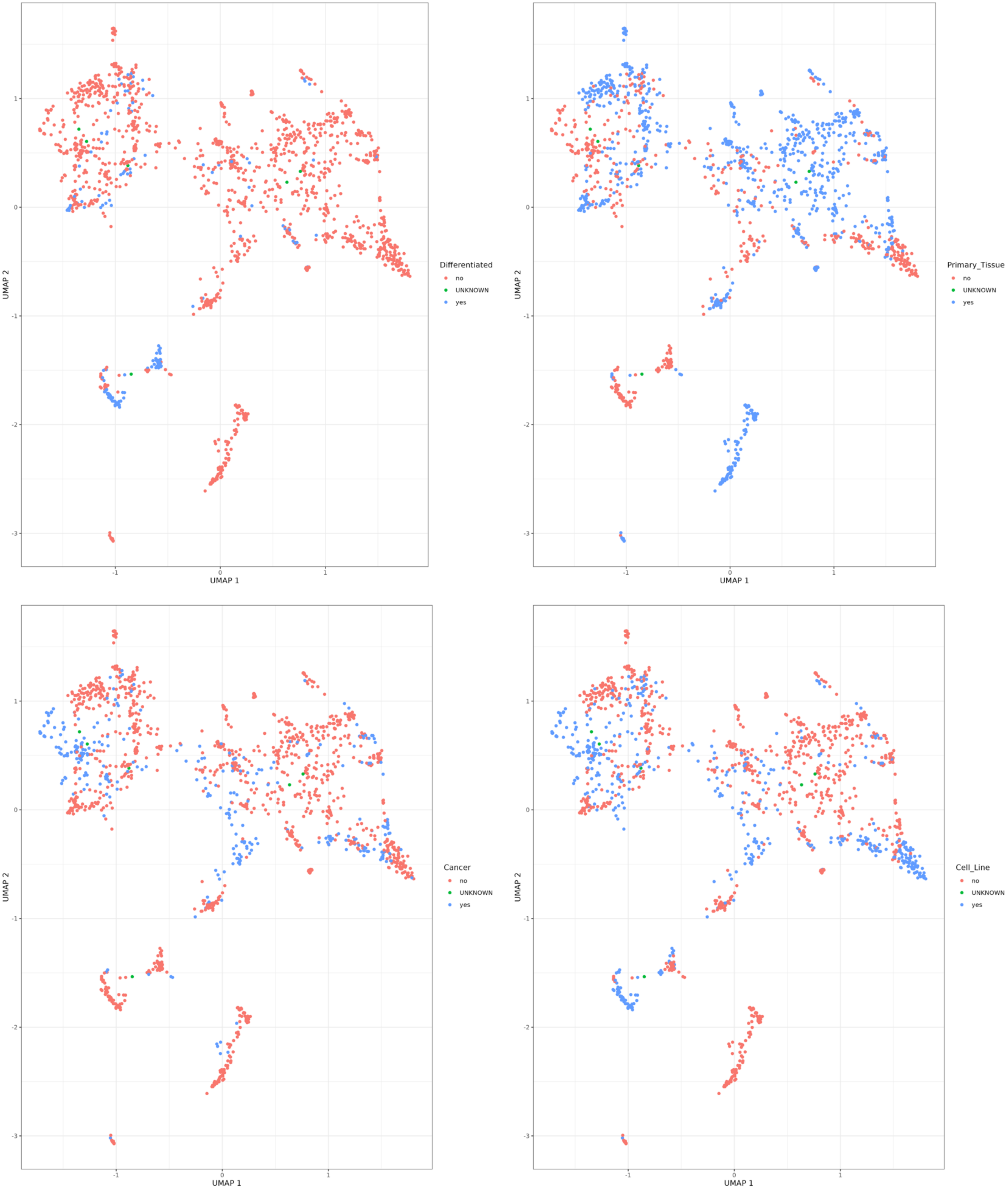


Supplementary Figure 18: Chromatin accessibility profile clustering results colored by sample metadata other than biological origin.

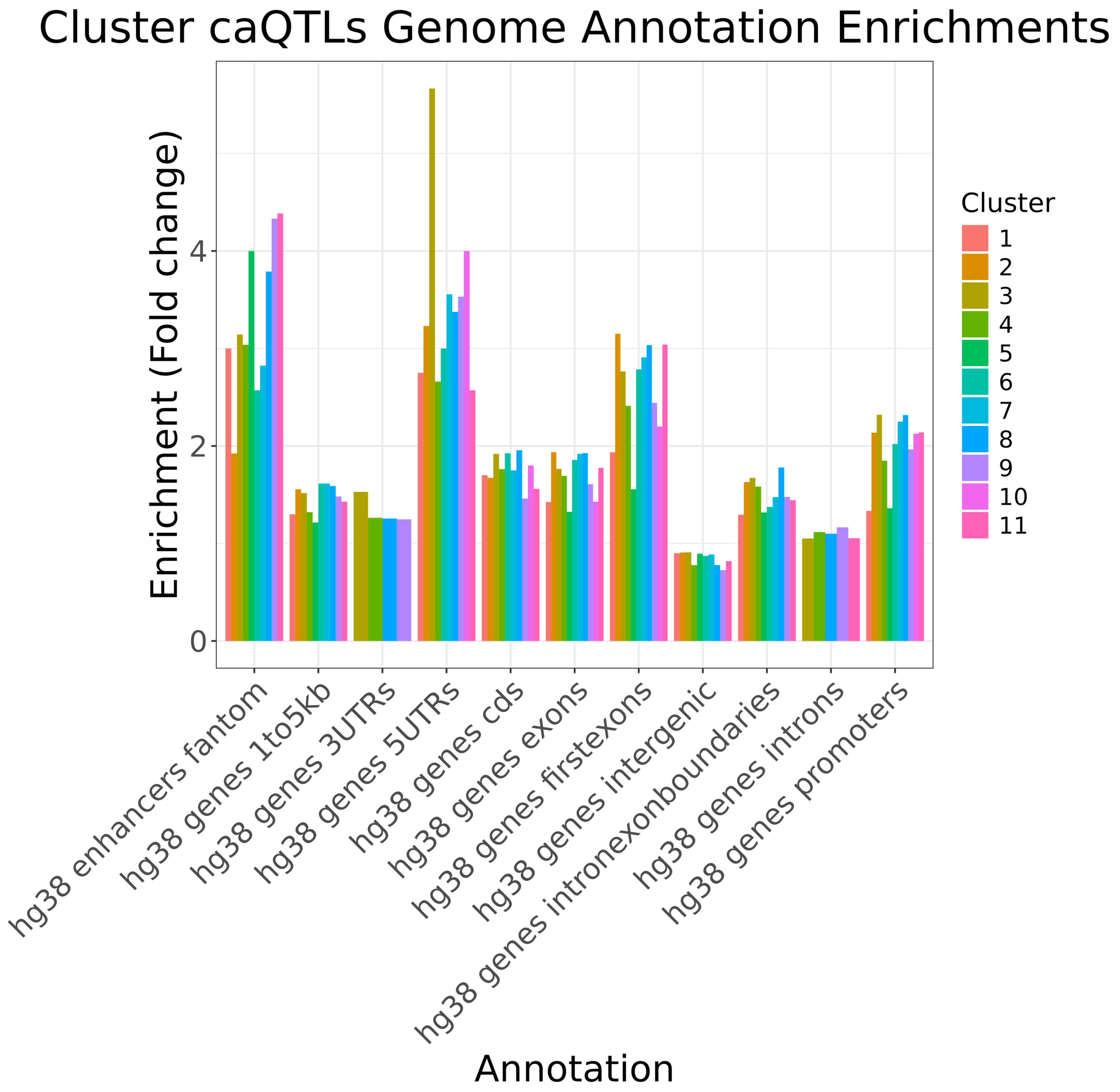


Supplementary Figure 19: Enrichment of caQTL peaks from each cluster, compared to matched random control regions, in various genomic annotation categories.

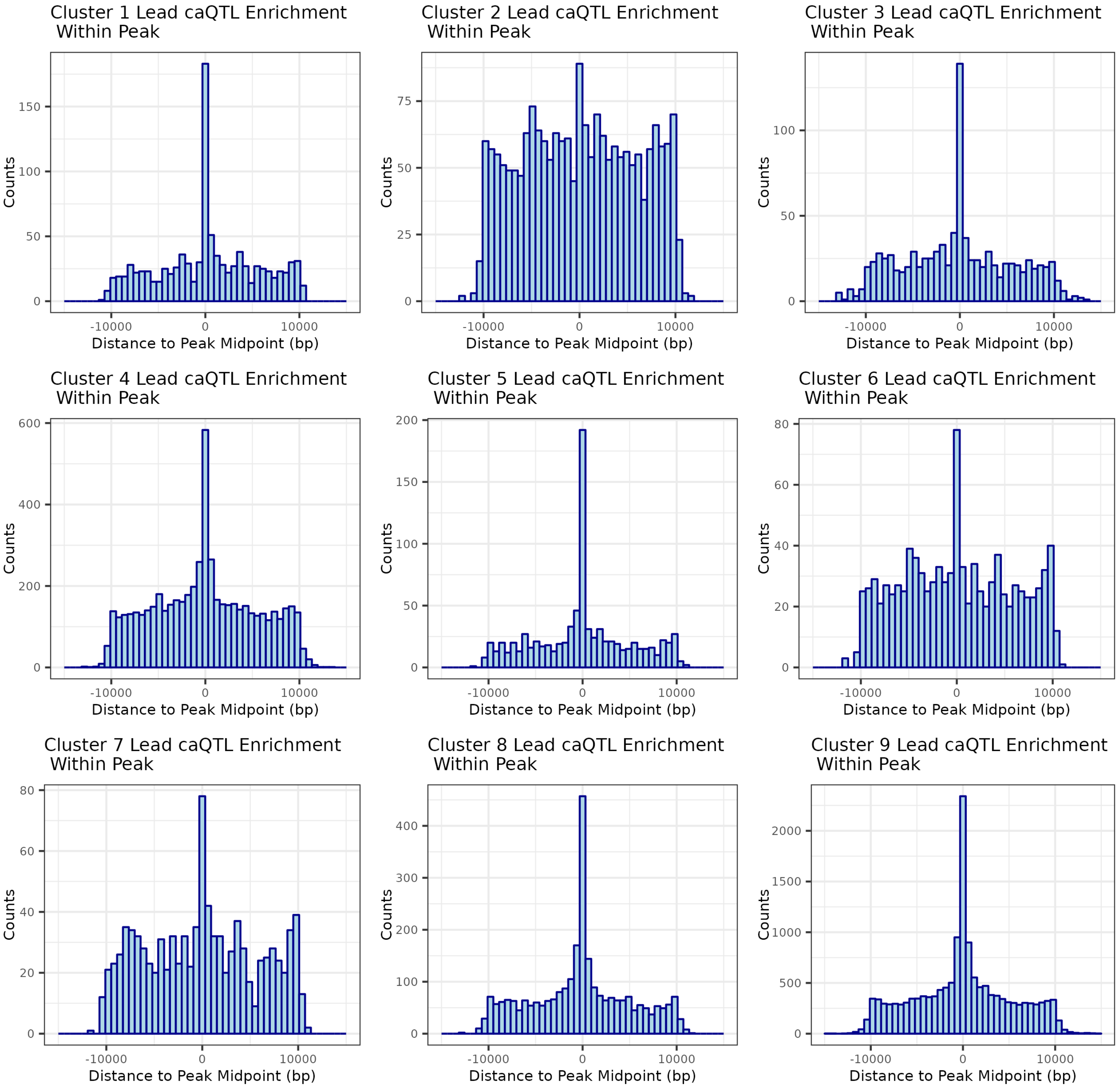


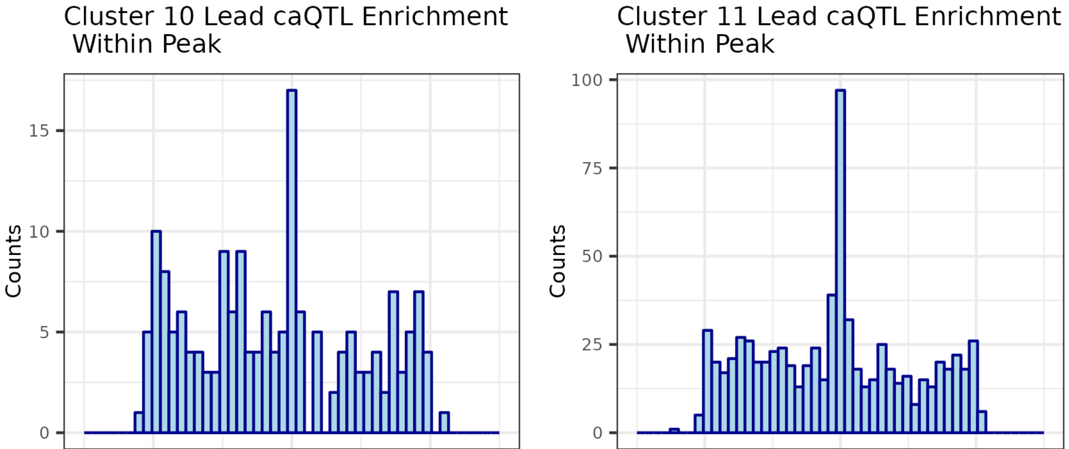


Supplementary Figure 20: Distance from cluster lead caQTL variant to midpoint of caQTL peak showing elevation of caQTL variant within the identified chromatin

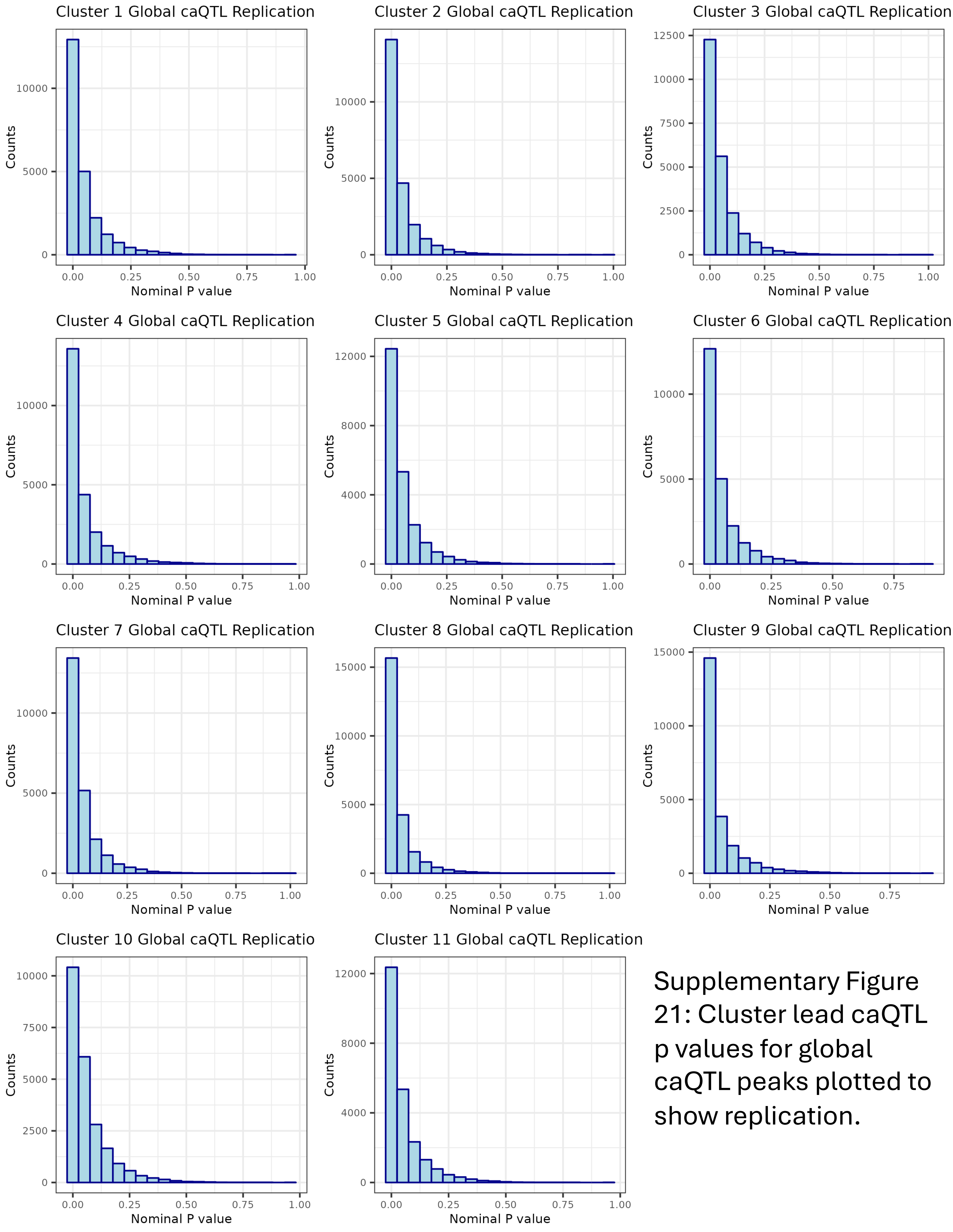


Supplementary Figure 21: Cluster lead caQTL p values for global caQTL peaks plotted to show replication.

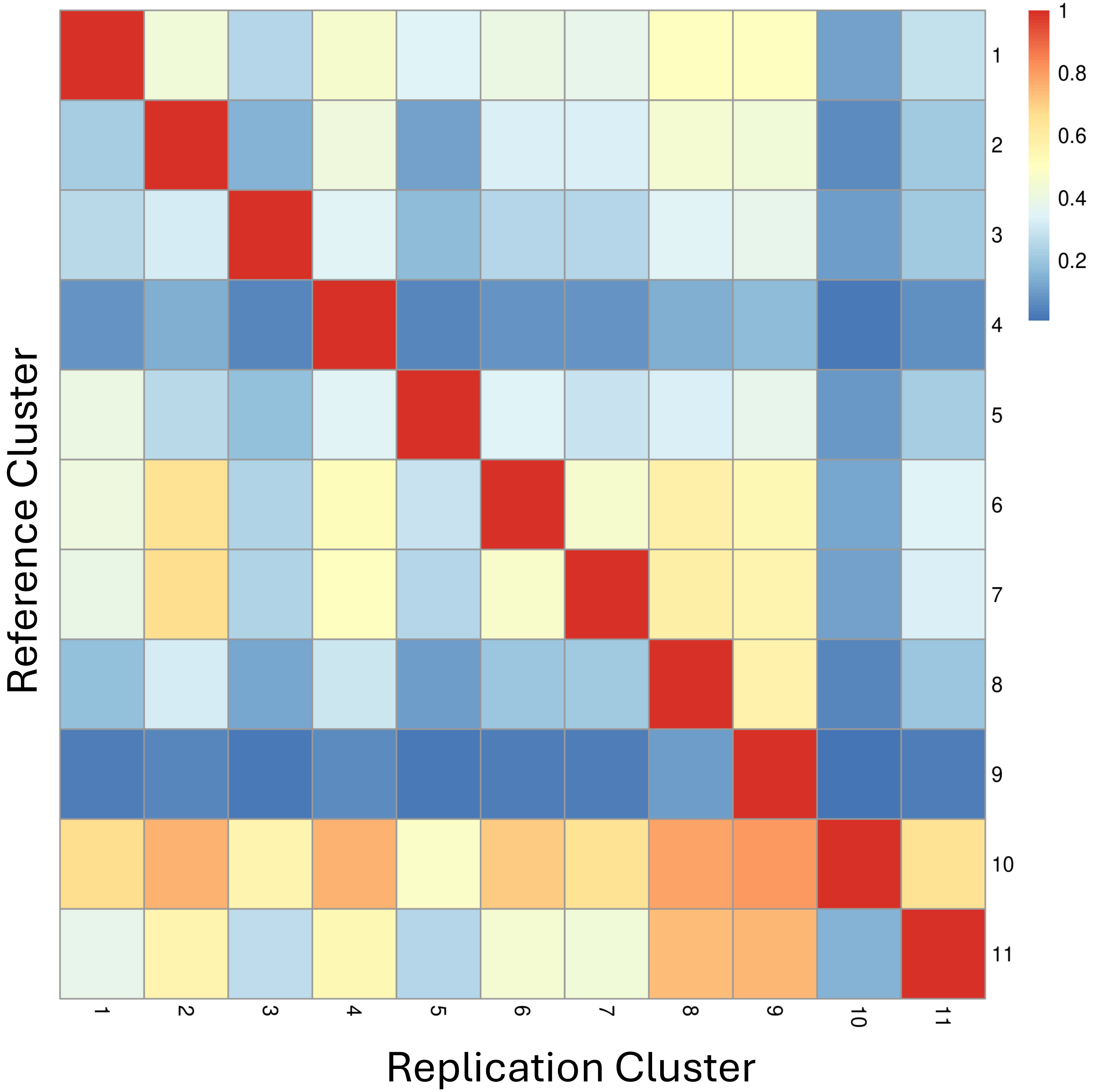


Supplementary Figure 22: Sharing of caQTL peaks across each cluster. Plotted across each row is the proportion of caQTL peaks, relative to the total number of reference cluster caQTL peaks, that are also identified as a caQTL peak in the replication cluster.

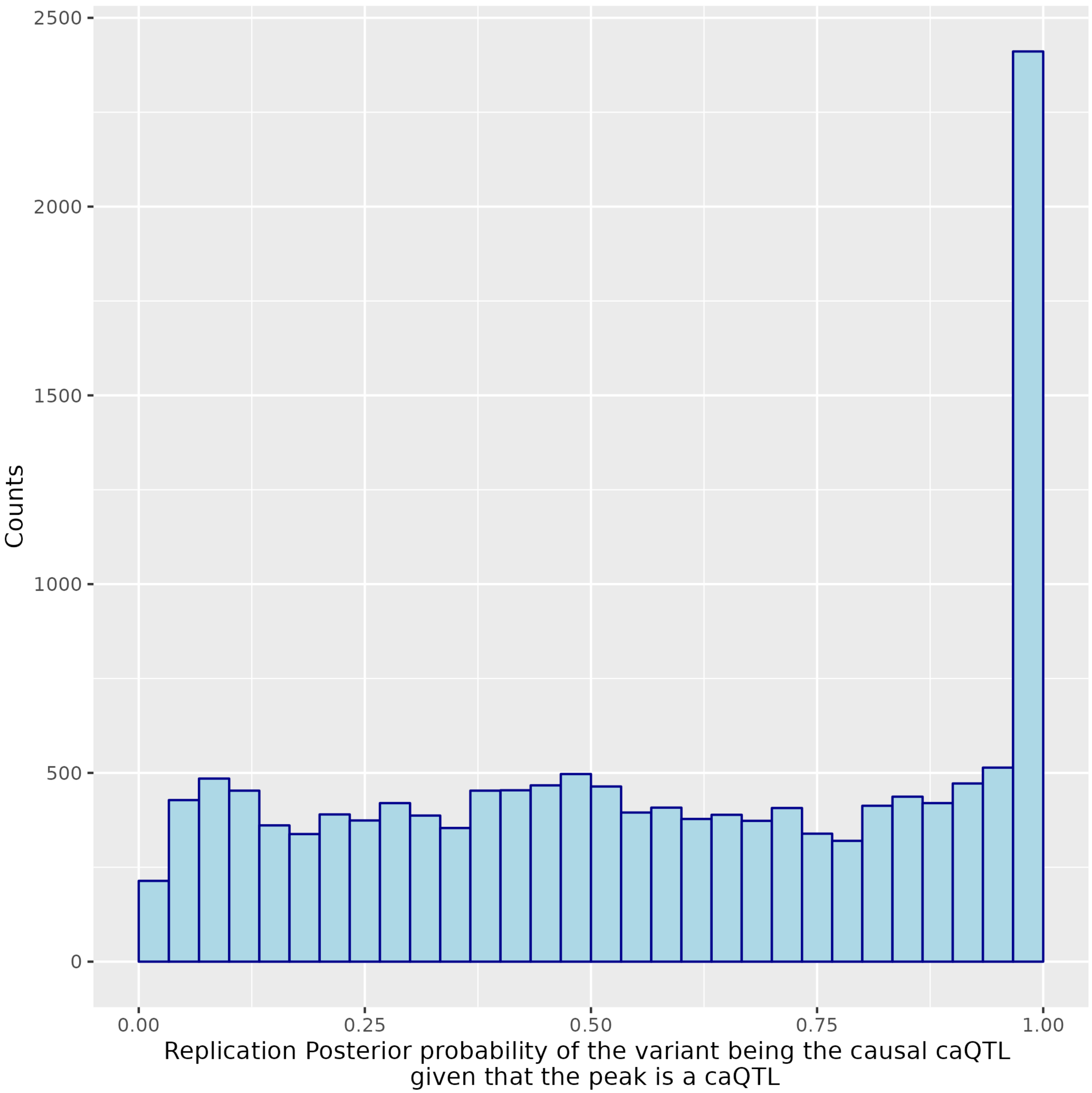


Supplementary Figure 23: Cluster 9 lead caQTL peaks matched to peaks from external dataset, from which 72 cluster 9 samples originated. Posterior probability of cluster 9 lead caQTL variant being causal for caQTL peak in exterior study plotted.

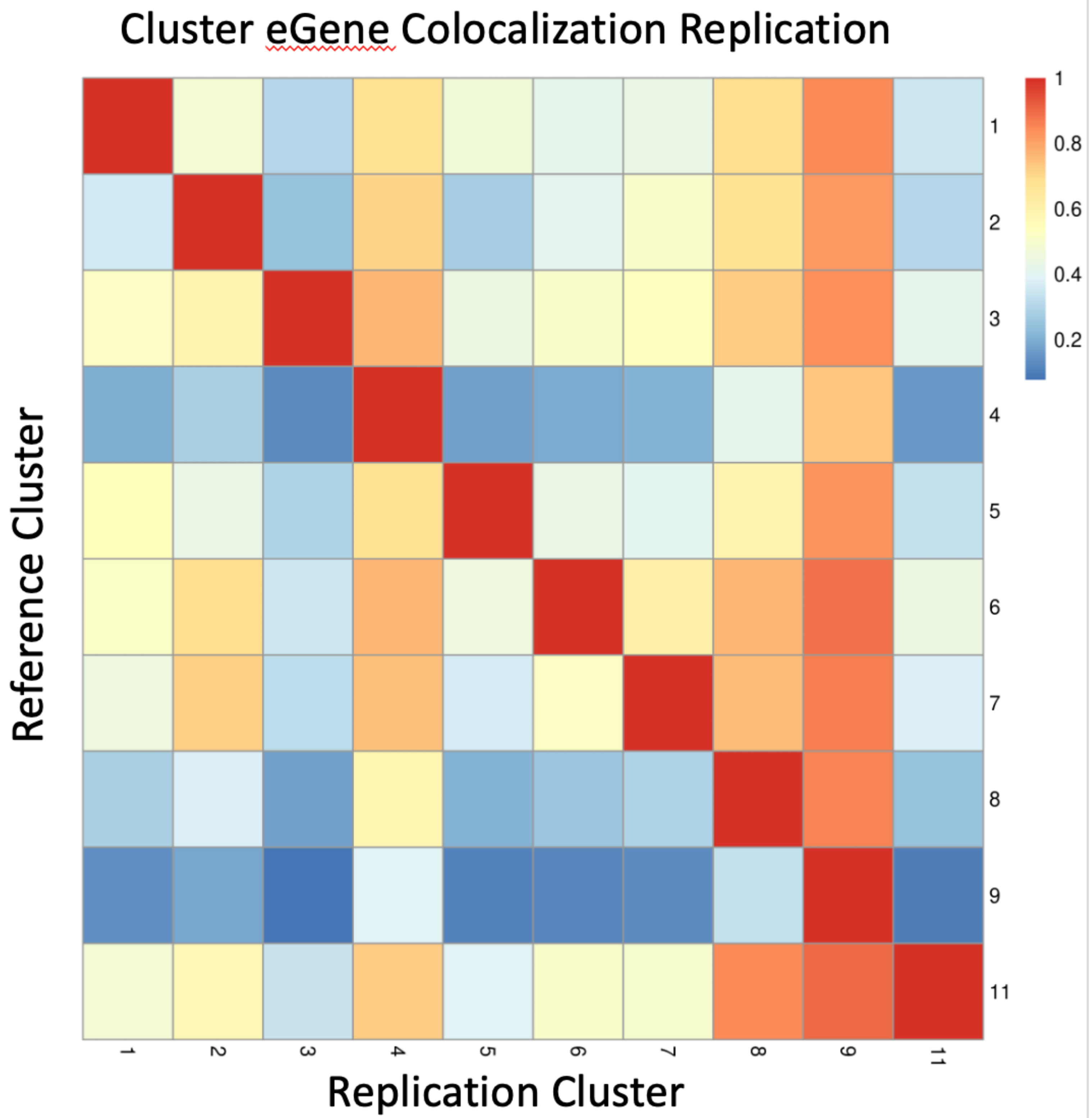


Supplementary Figure 24: Cluster caQTL and eQTL colocalization eGene sharing across clusters. Plotted is the proportion of total colocalizing eGenes in reference cluster, relative to the total number of reference cluster colocalizing eGenes, that also colocalized in the replication cluster.

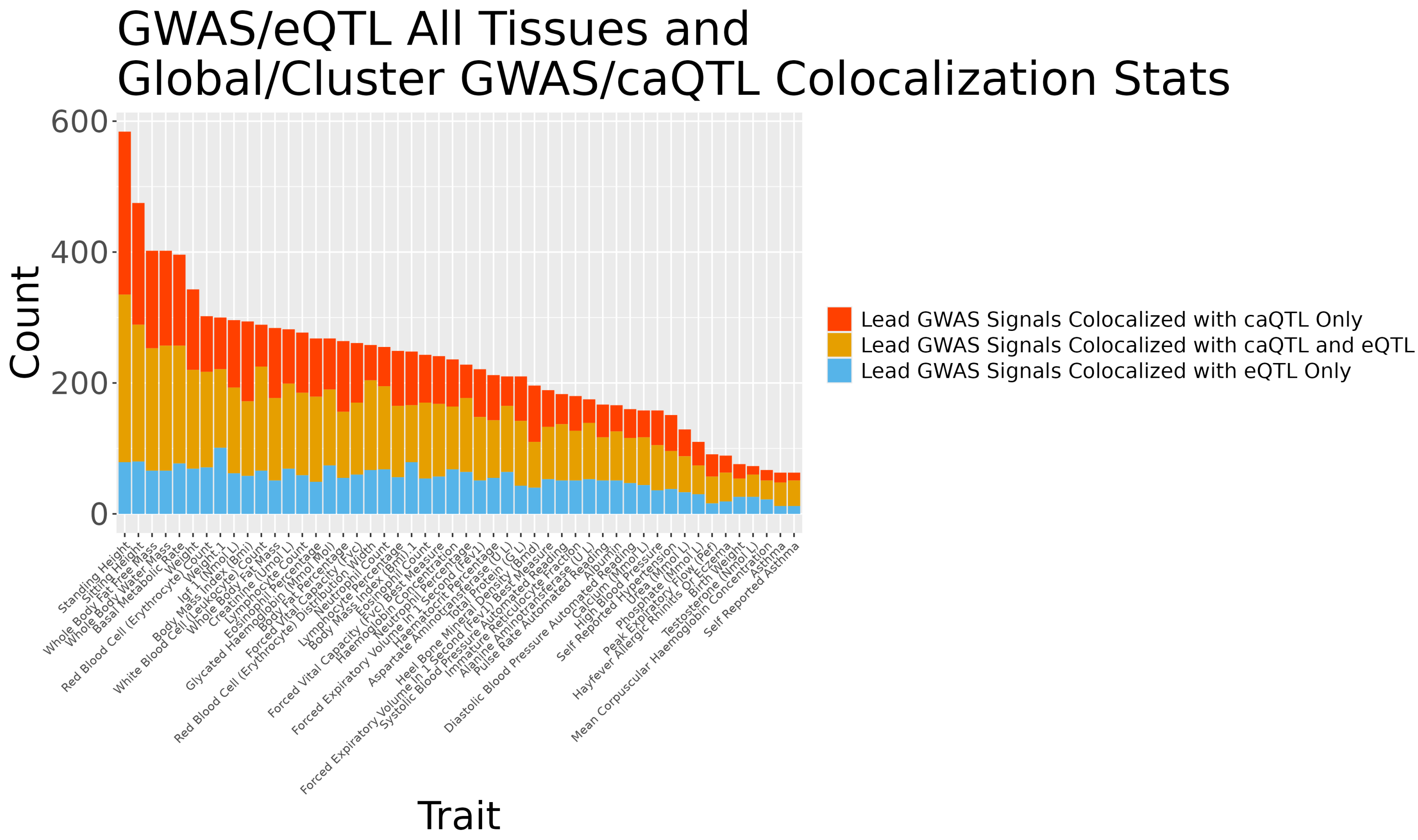


Supplementary Figure 25: For each GWAS trait, independent lead GWAS variant signals were checked for colocalization with global/cluster caQTL and eQTL signals across all GTEx tissues. Plotted is the number of unique lead GWAS signals per colocalization group, as multiple caQTL peaks, eGenes, etc. can colocalize with the same lead GWAS signal. Traits with greater than 50 colocalizing lead variants shown.

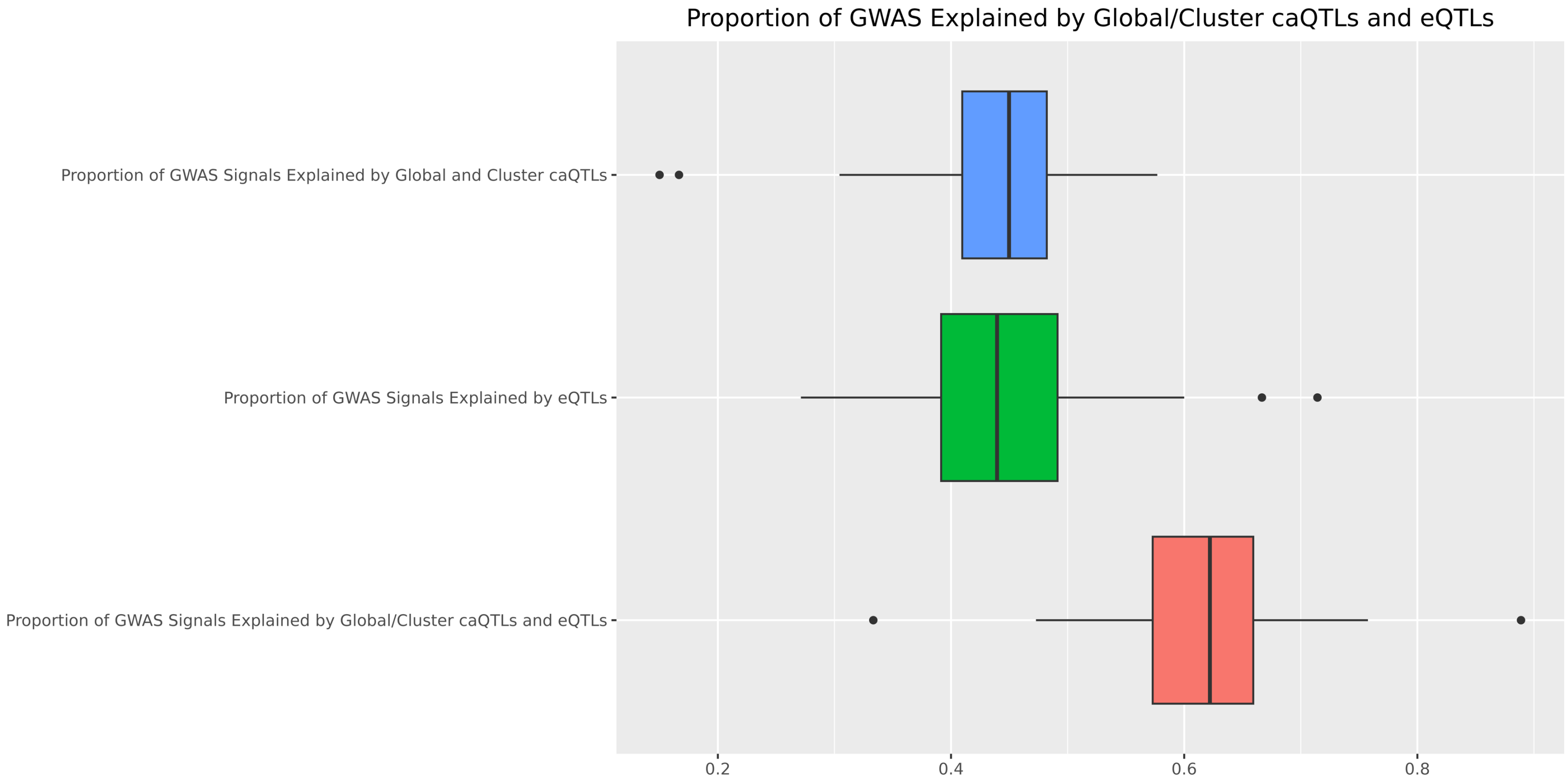


Supplementary Figure 26: Colocalizations were performed between global/cluster caQTL/GWAS and eQTL/GWAS. Each GWAS signal was checked to see if it colocalized exclusively with caQTLs, eQTLs, or colocalized with both. Proportion of tested GWAS signals that colocalized in each category are plotted.
